# Mitochondrial calcium influx-driven bioenergetics in the dopaminergic system selectively enable drug reward

**DOI:** 10.1101/2025.06.10.658191

**Authors:** Jun Gao, Haixin Zhao, Xiao Han, Lingmin Zeng, Guangqin Liu, Xuechen Wei, Changwei Liu, Wenjun Wu, Siqi Chen, Jiayi Chen, Ting Li, Jiye Yin, Tao Zhou, Xue-Min Zhang, Ai-Ling Li, Teng Li, Xin Pan

## Abstract

Drugs of abuse hijack the brain’s reward system, driving pathological dopamine surges that underlie compulsive behavior and addiction. However, directly targeting dopamine signaling for treatment risks disrupting natural reward processes. Here, we identify a bioenergetic mechanism that selectively promotes addiction-related dopamine release and behaviors. Opioids and methamphetamine, but not natural rewards, induce mitochondrial calcium (Ca^2+^) influx via the mitochondrial calcium uniporter (MCU) in dopaminergic terminals of the nucleus accumbens. Optogenetic stimulation reveals that this mitochondrial Ca^2+^ influx occurs exclusively during high-intensity dopaminergic neuronal activation. This Ca^2+^ influx drives rapid ATP production, compensating for energy deficits caused by neuronal hyperactivity and enabling sustained dopamine release. Genetic deletion or pharmacological inhibition of MCU in dopaminergic neurons selectively reduces drug-induced dopamine release and prevents addictive behaviors while sparing natural reward processing in mice. These findings uncover a distinct mitochondrial bioenergetic mechanism underlying drug reward and propose MCU as a promising therapeutic target for addiction treatment.

## Introduction

The brain’s reward system is a highly conserved network crucial for motivation and reinforcing survival behavior^1–3^. Central to this system is the mesolimbic circuit, particularly dopaminergic projections from the ventral tegmental area (VTA) to the nucleus accumbens (NAc), which play a critical role in reward processing. Natural rewards, such as eating or social interaction, trigger moderate dopamine release that induces pleasure and motivates these behaviors^4–6^. Addictive substances hijack this system, causing pathological dopamine elevation and intense euphoria, which initiate drug reward and positive reinforcement, further reinforcing drug-seeking behavior and solidifying addiction-related memories^7–10^. This dysregulation drives addiction, a chronic condition with severe health and social consequences^11^. Developing effective treatments for addiction is particularly challenging due to its complex neurobiological mechanisms, coupled with genetic, and psychological and social factors^12^. Directly targeting dopamine signaling, through approaches such as receptor knockout and the administration of receptor antagonists, has been shown to effectively suppress addictive behaviors in preclinical models^13,14^. However, this strategy inevitably disrupts natural reward processing and leads to adverse effects, including depression-like symptoms and motor disturbances^15–17^. Therefore, a better understanding of the mechanisms that differentiate the sustained dopamine release seen in addiction from the mild release during natural rewards is crucial. Such insights could provide a foundation for developing therapeutic strategies that selectively target addiction-related dopamine release without impairing normal reward processing.

The brain, as the most energy-demanding organ, relies on finely tuned energy production to sustain neural activities, including neurotransmitter release and synaptic function^18–20^. ATP is essential for processes such as synaptic vesicle cycling, ion gradient maintenance, and neurotransmitter loading^21–23^, with presynaptic terminals relying on specialized mitochondrial clusters to meet these demands^24^. These mitochondria play a crucial role in providing ATP for dopamine synthesis, packaging, and release^19,20,25^. Addictive drugs, such as opioids, trigger dopaminergic hyperactivation and intense dopamine release^26^, which impose substantial bioenergetic challenges on presynaptic terminals. Clinical imaging studies further reveal increased glucose utilization in reward-related brain regions of individuals with substance use disorders during both acute drug exposure and cue-induced craving states^27–29^. These findings reflect heightened energy demands throughout the addiction cycle. Understanding how these demands are met at the cellular level could offer novel insights into addiction mechanisms and provide new therapeutic strategies.

Ca^2+^ signaling plays a pivotal role in regulating neuronal functions, including excitability, synaptic transmission, and plasticity^30,31^. Mitochondrial Ca^2+^ uptake links cellular calcium signals to energy metabolism via the mitochondrial Ca^2+^ uniporter (MCU)^32,33^. Through MCU, cytosolic Ca^2+^ enter the mitochondria, enhancing the activity of enzymes in the tricarboxylic acid cycle and the electron transport chain, thereby driving ATP production^34,35^. Unlike baseline mitochondrial energy production, MCU-mediated energy supply is rapidly activated, particularly during high-energy demanding cellular processes, highlighting the specialized role of MCU in meeting acute energy demands^36,37^. In this study, we investigated mitochondrial Ca^2+^ dynamics in dopaminergic neurons under different rewarding conditions. Strikingly, we found that only drugs of abuse, but not natural rewards, triggered mitochondrial Ca^2+^ entry via MCU in dopaminergic terminals of the NAc. This mitochondrial Ca^2+^ influx provides essential energy support for addiction-related dopamine release and behaviors. These findings uncover a novel bioenergetic mechanism underlying addiction and highlight the potential of MCU as a therapeutic target for addiction treatment.

## Results

### Dopaminergic terminals in the NAc exhibit drug-specific mitochondrial calcium influx

The mesolimbocortical dopamine system plays a key role in regulating reward perception, motivation, and learning^38–40^. Dopaminergic projections from the ventral tegmental area (VTA) to the nucleus accumbens (NAc) are central to the brain’s reward circuitry^41^. To explore the potential involvement of mitochondrial Ca^2+^ in the reward system, we expressed a Cre-dependent mitochondrial Ca^2+^ indicator specifically in dopaminergic neurons by stereotaxically injecting AAV-DIO-4mt-jGCaMP8s into the VTA of *DAT-Cre* mice. Histological analysis confirmed that this mitochondria-localized Ca^2+^ indicator was expressed with ∼96% specificity in tyrosine hydroxylase positive (TH^+^) dopaminergic neurons (Fig. 1a and Extended Data Fig. 1a). Using implanted optic fiber recordings above the NAc, we monitored 4mt-jGCaMP8s signals in the NAc terminals of these mice in response to drug reward and natural reward (Fig. 1a). Fiber photometry revealed that single intraperitoneal (i.p.) injection of heroin (10 mg/kg), morphine (30 mg/kg), or methamphetamine (METH, 2 mg/kg) significantly increased the mitochondrial Ca^2+^ levels within dopaminergic terminals in the NAc, whereas cocaine (20 mg/kg) did not alter mitochondrial Ca^2+^ levels (Fig. 1b,c). Notably, heroin induced the most pronounced alteration. In contrast, food intake did not trigger significant changes in mitochondrial Ca^2+^ levels in the NAc when mice consumed small food pellets, with fluctuation levels similar to those observed with saline administration (Fig. 1b,c). These findings indicate that rewards from certain drugs, but not natural rewards, trigger a significant increase in mitochondrial Ca^2+^ influx in the NAc. To further visualize the mitochondrial Ca^2+^ dynamics, we employed a miniaturized two-photon microscope for *in vivo* imaging of Ca^2+^ levels in NAc dopaminergic terminals (Fig. 1d). Real-time imaging confirmed a significant increase in mitochondrial Ca^2+^ fluorescence in this region specifically after heroin administration, but not cocaine (Fig. 1e and Supplementary Video 1). Detailed analysis of regions of interest (ROIs) revealed that approximately 84% of mitochondria exhibited a rapid increase in Ca^2+^ levels following heroin administration (Fig. 1f–h). These findings collectively suggest that mitochondrial Ca^2+^ influx in dopaminergic terminals in the NAc was selectively responsive to certain addictive drugs, potentially due to distinct mechanisms of action that uniquely influence mitochondrial Ca^2+^ dynamics in this brain region.

**Fig. 1.**
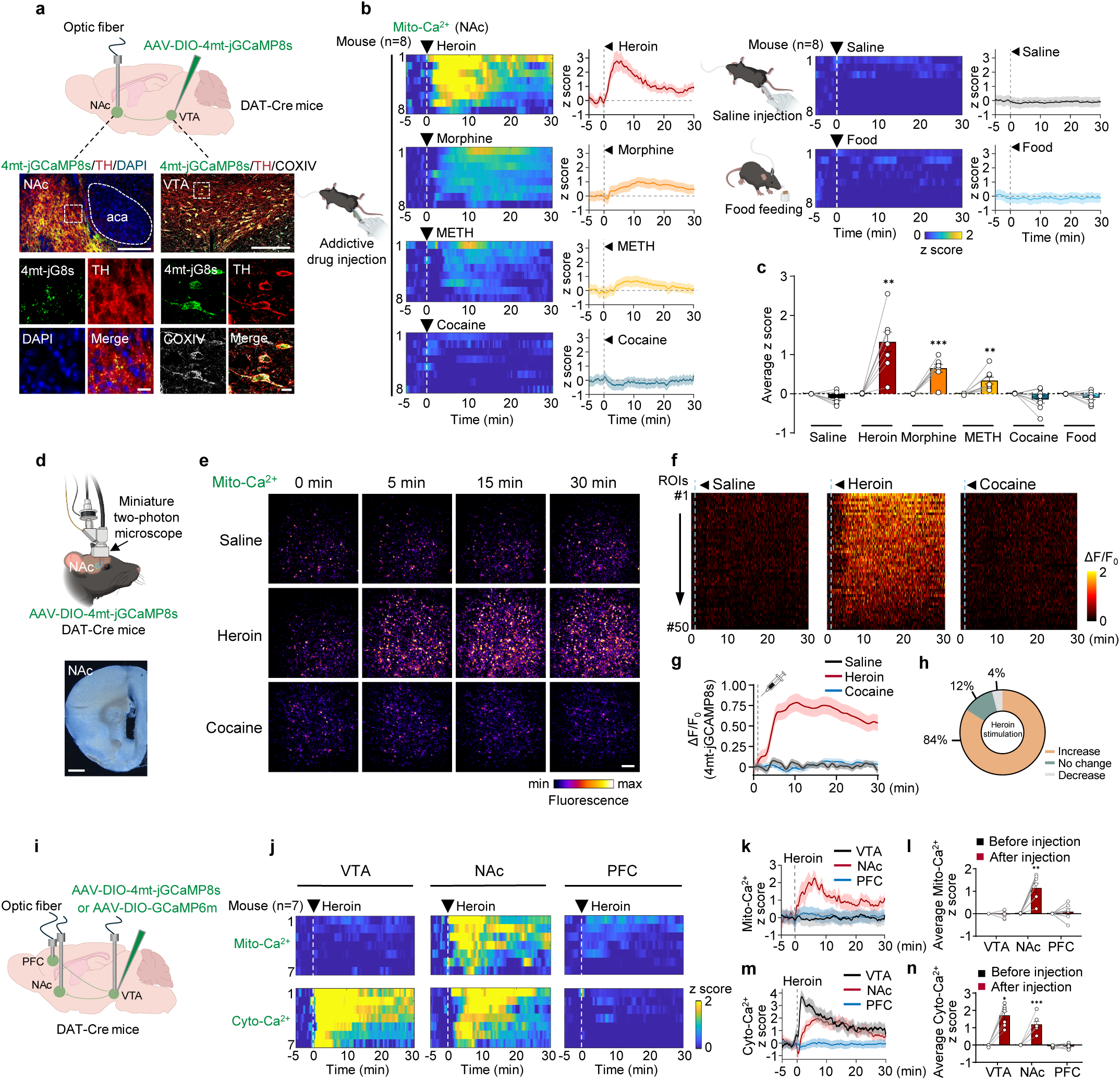
Dopaminergic terminals in the NAc exhibit drug-specific mitochondrial calcium influx. **a**, (Top) Schematic illustrating Cre-dependent AAV-DIO-4mt-jGCaMP8s expression in the VTA of DAT-Cre mice. Fiber photometry recorded mitochondrial Ca^2^⁺ signals from dopaminergic terminals in the NAc. (Bottom) Representative images showing 4mt-jGCaMP8s (4mt-jG8s, green) specifically expressed in tyrosine hydroxylase-immunopositive dopamine neurons (TH, red) in the NAc and VTA. Mitochondrial localization confirmed by COXIV staining (grey). Scale bars: 200 μm (top images); 20 μm (bottom enlarged images). **b**, Heatmaps (left) and traces (right) of mitochondrial Ca^2^⁺ dynamics (indicated by fluorescence of 4mt-GCaMP8s, z score) aligned to drug injection or food pellet consumption onset (white dotted line). Solid lines and shaded areas represent means ± SEM (*n* = 8 mice). **c**, Bar graphs showing the average z score of the 4mt-jGCaMP8s signals in the NAc 5 min before and 30 min after drug injection or food intake, corresponding to panel (**b**). **d**, (Top) Schematic of a fast high-resolution miniature two-photon microscope (FHIRM-TPM) mounted on a mouse’s head, used to record mitochondrial Ca^2+^ signals in the NAc. (Bottom) Image showing lens placement in the NAc. Scale bar: 1 mm. **e**, FHIRM-TPM time-lapse images showing enhanced mitochondrial Ca^2+^ influx after heroin administration compared to cocaine or saline. Scale bar: 20 μm. **f**, Heatmaps showing the changes in mitochondrial Ca^2+^ levels within 50 individual regions of interest (ROI) following intraperitoneal injections of saline, heroin (10 mg/kg), or cocaine (20 mg/kg). **g**, Traces of mitochondrial Ca^2^⁺ dynamics-indicated by fluorescence of 4mt-GCaMP8s, ΔF/F_0_) from panel (**f**) (n = 50 ROIs per group). **h**, Distribution of the percentage of ROIs displaying mitochondrial Ca^2^⁺ dynamics following heroin stimulation. **i**, Schematic of stereotaxic AAV injections (AAV-DIO-4mt-jGCaMP8s or AAV-DIO-GCaMP6m) into the VTA of DAT-Cre mice, with optic fibers placement in the VTA, NAc, or PFC for fiber photometry recordings. **j**, Heatmaps of mitochondrial Ca^2+^ (top) and cytoplasmic Ca^2+^ (bottom) fluorescence changes in the VTA (left), NAc (middle), and PFC (right), aligned to heroin injection onset (white dotted line, *n =* 7 mice). **k**, Traces of mitochondrial Ca^2^⁺ dynamics (indicated by 4mt-GCaMP8s fluorescence, z score) in the VTA (black), NAc (red), and PFC (blue) aligned to heroin injection (grey dotted line), corresponding to panel (**j**). **l**, Bar graph showing average z score of the 4mt-jGCaMP8s fluorescence signals in the VTA, NAc, and PFC 5 min before (black) and 30 min after (red) heroin injection, based on data from (**k**). **m**, Traces of cytoplasmic Ca^2^⁺ dynamics (indicated by GCaMP6m fluorescence, z score) in the VTA (black), NAc (red), and PFC (blue) before and after heroin injection, corresponding to panel (**j**). **n**, Bar graph showing average z score of the GCaMP6m fluorescence signals in the VTA, NAc, and PFC 5 min before (black) and 30 min after (red) heroin injection, based on data from (**m**). All data are presented as means ± SEM. Statistical analyses include paired two-tailed t-test for comparing pre-and post-reward (**c**[saline, heroin, and morphine], **l**[VTA, NAc], **n**[NAc, PFC]) or Wilcoxon matched-pairs signed rank tests (**c**[METH, cocaine, and food], **l**[PFC], **n**[VTA]). \**P* < 0.05, \*\**P* < 0.01, \*\*\**P* < 0.001.

### Regional specificity of drug-induced mitochondrial calcium influx

Dopaminergic projections from the VTA to the NAc and prefrontal cortex (PFC) are crucial for the neural circuitry underlying drug addiction^42^. To investigate the regional specificity of mitochondrial Ca^2+^ influx in response to heroin in these brain regions, we positioned optical fibers above 4mt-jGCaMP8s-expressing dopaminergic neurons in the VTA or their terminals in the NAc and PFC (Fig. 1i and Extended Data Fig. 1b). Interestingly, only dopaminergic terminals in the NAc exhibited a significant increase in mitochondrial Ca^2+^ levels upon heroin injection (Fig. 1j–l). In contrast, minimal changes in mitochondrial Ca^2+^ levels were observed in the dopaminergic somas of the VTA and terminals in the PFC (Fig. 1j–l). Given that mitochondrial Ca^2+^ uptake depends on cytosolic Ca^2+^ elevation^43,44^, we investigated whether the observed mitochondrial Ca^2+^ changes were associated with cytosolic Ca^2+^ elevation. To this end, we expressed a Cre-dependent cytosolic Ca^2+^ indicator (GCaMP6m) in VTA dopaminergic neurons of *DAT-Cre* mice (Fig. 1i). Histological analysis confirmed ∼91% specificity of this cytosolic Ca^2+^ indicator for TH^+^ dopaminergic neurons (Extended Data Fig. 1c,d). Intriguingly, heroin administration induced a marked elevation of cytosolic Ca^2+^ within both VTA somas and NAc terminals, but not in PFC terminals (Fig. 1j,m,n). Thus, while heroin triggered cytosolic Ca^2+^ increase in both VTA somas and NAc terminals, mitochondrial Ca^2+^ influx was observed only in NAc terminals (Fig. 1j–n). Overall, these results suggest a region-specific mechanism where opioid drugs trigger robust mitochondrial Ca^2+^ influx specifically at dopamine projection terminals within the NAc, potentially playing a unique role in regulating drug addiction.

### MCU-mediated mitochondrial calcium influx is essential for drug-induced reward behaviors in mice

Mitochondrial calcium influx is mediated by the MCU located in the inner mitochondrial membrane^45,46^. To investigate its role in drug-induced reward behavior, we generated dopaminergic neuron-specific MCU knockout mice (*DAT-Cre; MCU^flox/flox^*) by crossing DAT-Cre mice with *MCU* floxed mice (Extended Data Fig. 2a). Immunofluorescence staining confirmed efficient and specific MCU knockout in dopaminergic neurons (Fig. 2a). Using fiber photometry, we observed that heroin-induced enhancement of mitochondrial Ca^2+^ fluorescence in NAc dopaminergic terminals was significantly suppressed in *DAT-Cre; MCU^flox/flox^* (DAT-*MCU* cKO) mice compared to *DAT-Cre; MCU^+/+^* (wild-type, WT) controls (Fig. 2b,c), while cytosolic Ca^2+^ elevation in VTA somas remained unaffected by MCU depletion (Extended Data Fig 2b,c). These results indicate that heroin-induced mitochondrial Ca^2+^ influx in dopaminergic terminals is mediated by MCU.

**Fig. 2.**
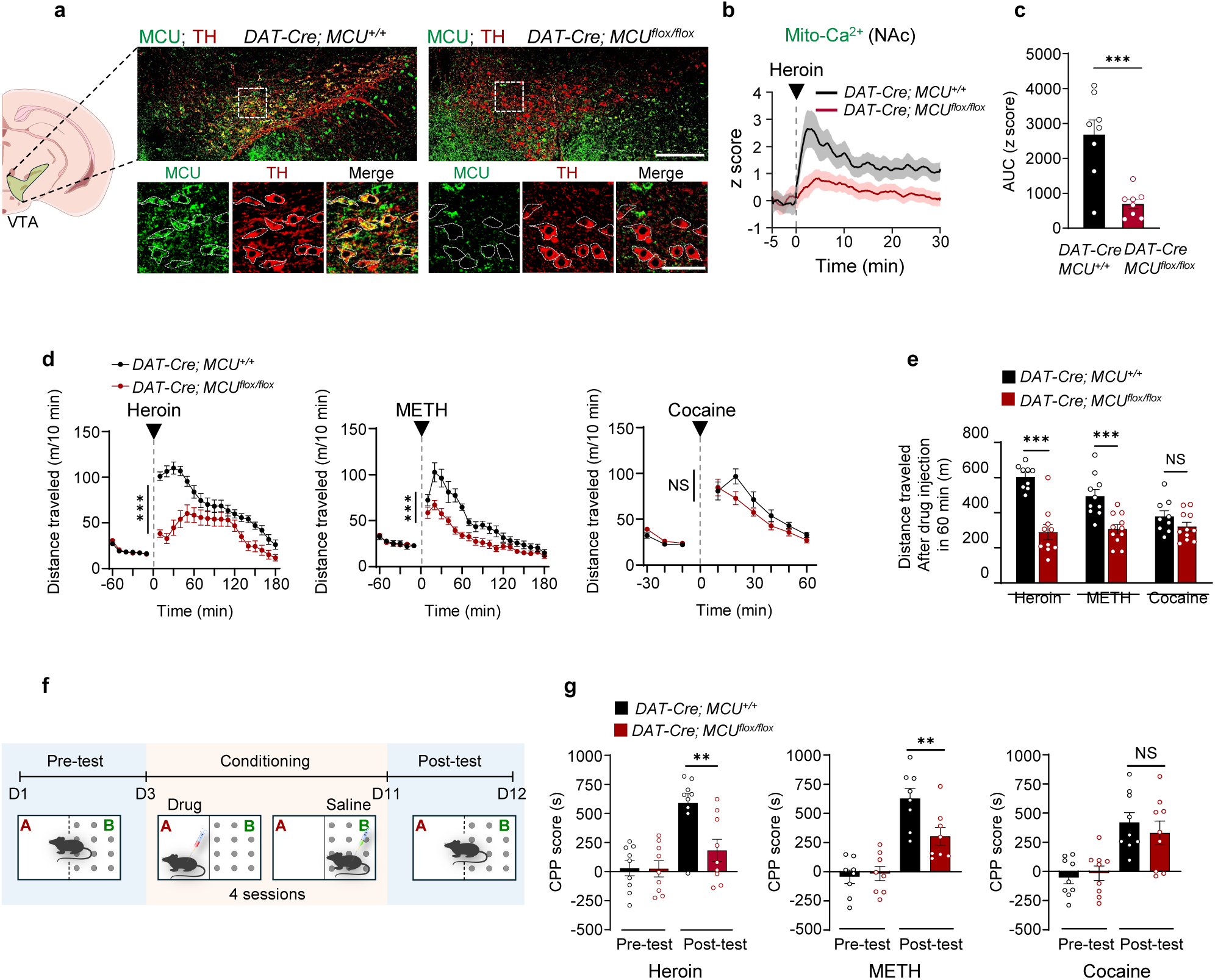
MCU-mediated mitochondrial calcium influx is essential for drug reward behaviors in mice. **a**, Representative immunofluorescent images showing MCU (green) and TH (red) expression in the VTA of *DAT-Cre; MCU^+/+^*(WT) and *DAT-Cre; MCU^flox/flox^*(DAT-*MCU* cKO) mice. Scale bar: 200 μm (upper); 50 μm (lower). **b**, Traces of mitochondrial Ca^2^⁺ dynamics (4mt-GCaMP8s fluorescence, z score) in NAc dopaminergic terminals aligned to heroin injection (grey dotted line and arrow) in WT and DAT-*MCU* cKO mice (*n =* 8 per group). **c**, Bar graph showing area under the curve (AUC) of mitochondrial Ca^2+^ signals (4mt-GCaMP8s fluorescence) in the NAc during the 30-min period after heroin injection, based on data from (**b**). **d**, Distance traveled in open field for WT (black) and DAT-*MCU* cKO (red) mice before and after heroin (left, 10 mg/kg), methamphetamine (METH, middle, 2 mg/kg), or cocaine (right, 20 mg/kg) injection. Grey dotted lines and black arrowheads mark injection times. Sample sizes: heroin (WT: *n* = 9; DAT-*MCU* cKO: *n* = 10), METH (WT: *n* = 10; DAT-*MCU* cKO: *n* = 11), and cocaine (WT: *n* = 9; DAT-*MCU* cKO: *n* = 11). **e**, Cumulative distance traveled within 60 minutes after heroin, METH, or cocaine injection, derived from locomotor activity data in (**d**). **f**, Experimental timeline for drug-conditioned place preference (CPP). **g**, CPP scores during pre-test and post-test sessions for heroin (*n =* 9 per group), METH (*n =* 8 per group), and cocaine (*n =* 9 per group) in WT and DAT-*MCU* cKO mice. Data are presented as means ± SEM. Statistical analyses included two-sided unpaired Student’s t-test (**c**, **e**) and two-way ANOVA with Bonferroni post hoc test (**d**, **g**). NS, not significant; \**P* < 0.05, \*\**P* < 0.01, \*\*\**P* < 0.001.

As previously demonstrated above, different drugs showed distinct patterns of mitochondrial Ca^2+^ responses, with opioids and METH inducing significant increases while cocaine showed minimal effects (Fig. 1b,c). We next employed two established assays: the locomotor activity assay and conditioned place preference (CPP) test to assess the functional impact of MCU on drug reward-related behaviors. Addictive drugs, including heroin (10 mg/kg), morphine (30 mg/kg), or methamphetamine (METH, 2 mg/kg) induced robust hyperlocomotion in WT mice in open-field tests. In contrast, DAT-*MCU* cKO mice exhibited significantly reduced locomotor activity following drug administration (Fig. 2d,e and Extended Data Fig. 2d). Interestingly, cocaine-induced locomotion (20 mg/kg) remained unaffected by MCU deletion (Fig. 2d,e). These findings suggest that MCU is essential for the locomotor activity induced by opioids and METH, but not cocaine. We further investigated the role of MCU in reward learning using the CPP test (Fig. 2f). Following conditioning, WT mice exhibited a significant preference for the drug-paired compartment, spending substantially more time in this area. In contrast, DAT-*MCU* cKO mice spent significantly less time than WT control mice in the compartment previously paired with heroin or METH (Fig. 2g), indicating that MCU deficiency impairs reward learning associated with these drugs. However, cocaine-induced CPP remained unaltered in DAT-*MCU* cKO mice (Fig. 2g). These results, aligned with our observations of mitochondrial Ca^2+^ responses under drug stimuli (Fig. 1b,c), support the notion that MCU-mediated mitochondrial Ca^2+^ influx in NAc dopaminergic terminals is crucial for drug reward-related behaviors induced by opioids and METH, but not cocaine.

To validate these findings through an independent approach, we employed viral-mediated conditional knockout using AAV expressing Cre recombinase under the control of the tyrosine hydroxylase (TH) promoter. Stereotaxic injection of AAV-TH-Cre into the VTA of *MCU* floxed mice (*MCU^flox/flox^*) mice generated dopamine neuron-specific MCU knockout mice (AAV-*TH-Cre; MCU^flox/flox^*). Immunofluorescence staining confirmed efficient and specific MCU knockout in dopaminergic neurons (Extended Data Fig. 2e). In agreement with our findings using *DAT-Cre; MCU^flox/flox^* mice, AAV-*TH-Cre; MCU^flox/flox^*mice displayed comparable reductions in heroin-induced locomotor activity and CPP compared to controls, with no significant change in cocaine-induced behaviors (Extended Data Fig. 2f–h). Locomotor sensitization assays also indicated that MCU deficiency impaired heroin-induced sensitization but not cocaine-induced sensitization (Extended Data Fig. 2i,j). These results further solidify the crucial role of MCU in dopaminergic terminals for drug reward-related behaviors induced by opioid drugs, but not cocaine.

### Mitochondrial calcium influx is required for heroin and METH-induced dopamine elevation in the NAc

Dopamine levels in the NAc are essential for reward processing, and all known addictive drugs elevate dopamine levels in this region^9,41^. Given the above results that MCU-mediated mitochondrial Ca^2+^ influx is integral to drug-induced reward behaviors, we further investigated its role in regulating dopamine levels in the NAc. To simultaneously record both dopamine and mitochondrial Ca^2+^ dynamics in the NAc, we expressed a red fluorescent dopamine sensor (rDA1m^47^) in the NAc and the mitochondrial calcium indicator (4mt-jGCaMP8s) in the VTA of DAT-Cre mice, and implanted optic fiber above the NAc (Fig. 3a,b). Fiber photometry recordings revealed that heroin administration triggered synchronized increase in both dopamine and mitochondrial Ca^2+^ signals (Fig. 3c). When these mice were ranked according to their increase in mitochondrial Ca^2+^ levels following heroin stimulation, they exhibited similar patterns in the elevation of dopamine levels in the NAc (Fig. 3d). A further correlation analysis demonstrated that the amplitudes of dopamine increase were positively correlated with that of mitochondrial Ca^2+^ in the NAc of these mice (Fig. 3e).

**Fig. 3.**
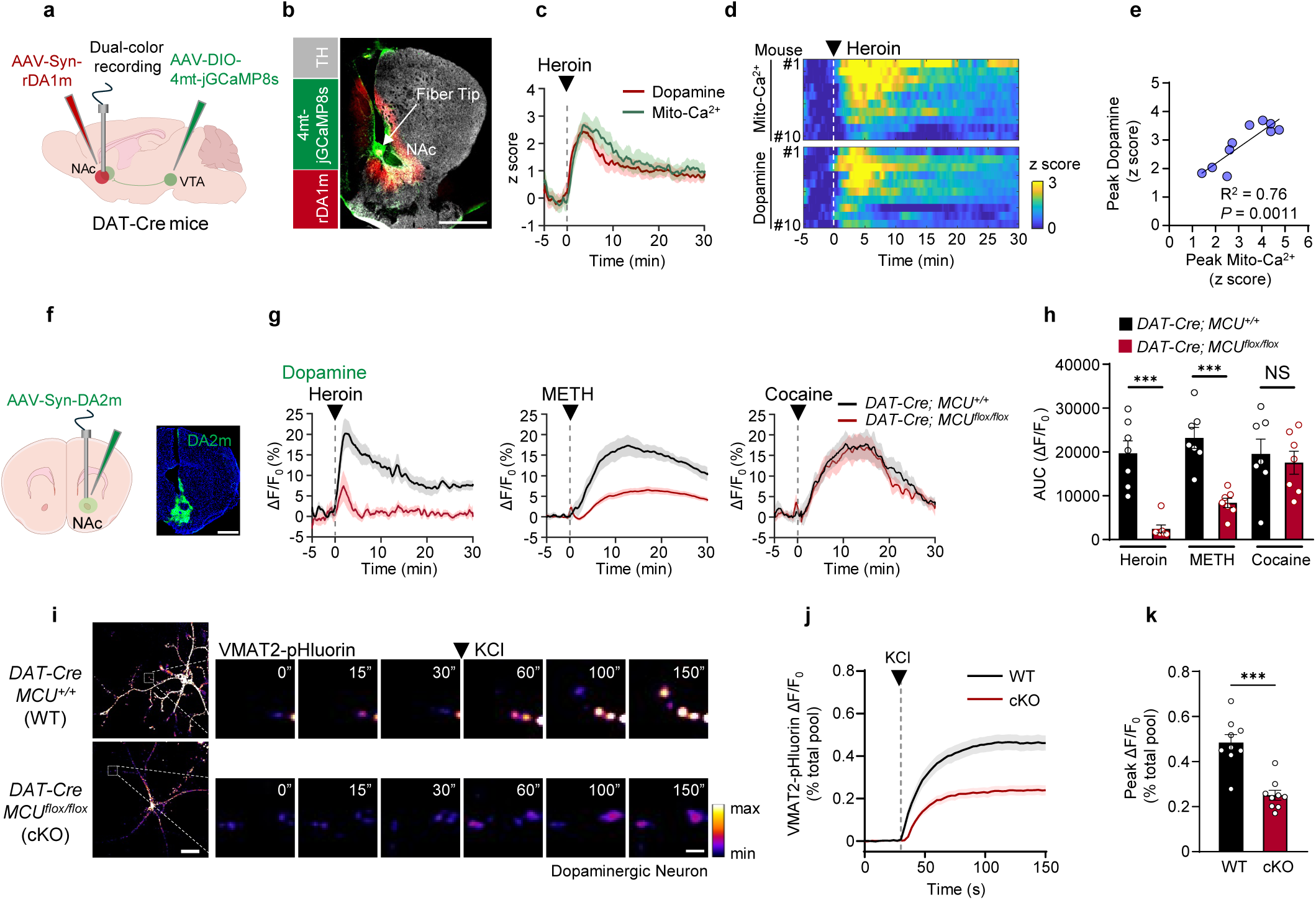
Mitochondrial calcium influx is required for dopamine release. **a**, Schematic showing unilateral viral delivery of AAV-DIO-4mt-jGCaMP8s (green) in the VTA and AAV-Syn-rDA1m (red) in the NAc of DAT-Cre mice, with an optic fiber implanted in the NAc for dual-color recording of mitochondrial Ca^2+^ and dopamine signals. **b**, Representative image displaying 4mt-jGCaMP8s (green) in TH^+^ neurons (grey) and Syn-rDA1m (red) in the NAc. Scale bar: 1 mm. **c**, Traces of dopamine signals (indicated by rDA1m fluorescence, z score, red) and mitochondrial Ca^2+^ dynamics (indicated by 4mt-jGCaMP8s fluorescence, z score, green) in the NAc aligned to heroin injection (black arrowhead and grey dotted line) (*n* = 10 mice). **d**, Heatmaps of mitochondrial Ca^2+^ dynamics (top) and dopamine signals (bottom) aligned to heroin injection onset, with each row representing paired recordings from the same mouse (*n* = 10 mice). **e**, Pearson correlation analysis of peak z-scores for dopamine and mitochondrial Ca^2+^ signals in the NAc over 30 minutes after heroin injection (R^2^ = 0.76, \*\**P* = 0.0011), corresponding to panel (**d**). **f**, (Left) Schematic showing AAV-Syn-DA2m delivery to the NAc and fiber photometry recording location. (Right) Representative image of DA2m expression in the NAc. Scale bar: 1 mm. **g**, Traces of dopamine signals (indicated by DA2m fluorescence, ΔF/F_0_) in the NAc from *DAT-Cre; MCU^+/+^* (black) and *DAT-Cre; MCU^flox/flox^* (red) mice after heroin (left, 10 mg/kg), methamphetamine (METH, middle, 2 mg/kg), or cocaine (right, 20 mg/kg) injections (*n =* 7 mice per group). **h**, Bar graphs showing AUC of dopamine fluorescence signals (ΔF/F_0_) in the NAc during the 30-min period following heroin, METH, or cocaine injection in *DAT-Cre; MCU^+/+^* (black) and *DAT-Cre; MCU^flox/flox^* (red) mice, based on data from (**g**). **i**, Representative time-lapse images of VMAT2-pHluorin fluorescence in dopaminergic neurons after KCl stimulation. Scale bar: 10 μm (left), 1 μm (right) . **j**, Traces of VMAT2-pHluorin fluorescence in dopaminergic neurons following KCl stimulation, normalized to maximal fluorescence in an alkaline buffer (representing the total pool), as shown in (**i**). (*n =* 9 cells per group). **k**, Bar graphs showing the normalized peak fluorescence of VMAT2-pHluorin, representing dopamine exocytosis after KCl stimulation, based on data from (**j**). Data are presented as means ± SEM. Statistical analysis included Pearson correlation test (**e**), two-sided unpaired Student’s t-test (**h**[METH, cocaine], **k**), and Mann-Whitney test (**h**[heroin]). NS, not significant; \**P* < 0.05, \*\**P* < 0.01, \*\*\**P* < 0.001.

To further elucidate the causal relationship between mitochondrial Ca^2+^ influx and dopamine elevation, we compared drug-induced dopamine elevation in the NAc of WT and DAT-*MCU* cKO mice, using a green fluorescent dopamine sensor (DA2m^47^) (Fig. 3f). Heroin and METH both induced a rapid increase in dopamine levels in the NAc. However, this dopamine surge was significantly suppressed in the NAc of DAT-*MCU* cKO mice (Fig. 3g,h). In contrast, cocaine-induced dopamine elevation remained unaffected by MCU deletion (Fig. 3g,h). These findings suggest that MCU-mediated mitochondrial Ca^2+^ influx regulates opioid and METH reward by facilitating dopamine elevation in the NAc.

Among the addictive drugs we tested, only those known to stimulate dopamine release, such as opioids (heroin and morphine)^10^ and METH^48^, induced this robust influx of mitochondrial Ca^2+^. This observation suggests that mitochondrial Ca^2+^ influx may directly regulate dopamine release. To explore this possibility, we monitored dopamine release rate using VMAT2-pHluorin indicator specifically expressed in primary dopaminergic neurons isolated from WT and DAT-*MCU* cKO mice (Extended Data Fig. 3a–e). Upon KCl-induced depolarization of dopaminergic neurons, a significant increase in dopamine exocytosis, as indicated by the increased VMAT2-pHluorin fluorescent intensity, was observed in WT neurons. In contrast, this effect was markedly inhibited in MCU-deficient neurons (Fig. 3i-k). These results collectively suggest that MCU is required for dopamine release.

### MCU facilitates dopamine release by promoting ATP production

We next sought to explore the molecular mechanism by which MCU facilitates dopamine release. We hypothesized that the MCU-mediated mitochondrial Ca^2+^ influx, a process known to enhance mitochondrial energy production, provided the necessary energy for sustained dopamine release, particularly during robust neuronal activation induced by addictive drugs. To test this hypothesis, we employed the mitochondrial ATP probe 4mt-ATeam1.03 and red mitochondrial Ca^2+^ indicator 4mt-RCaMPh to simultaneously monitor mitochondrial ATP and Ca^2+^ dynamics in dopaminergic neurons. Following KCl-induced depolarization and neuronal activation, primary dopaminergic neurons exhibited a rapid increase in mitochondrial Ca^2+^ levels, which was accompanied by a marked rise in mitochondrial ATP production. However, this response was absent in neurons derived from DAT-MCU cKO mice (Fig. 4a,b). These results suggest that MCU-mediated mitochondrial Ca^2+^ influx is essential for promoting mitochondrial ATP production during dopaminergic neuronal activation.

**Fig. 4.**
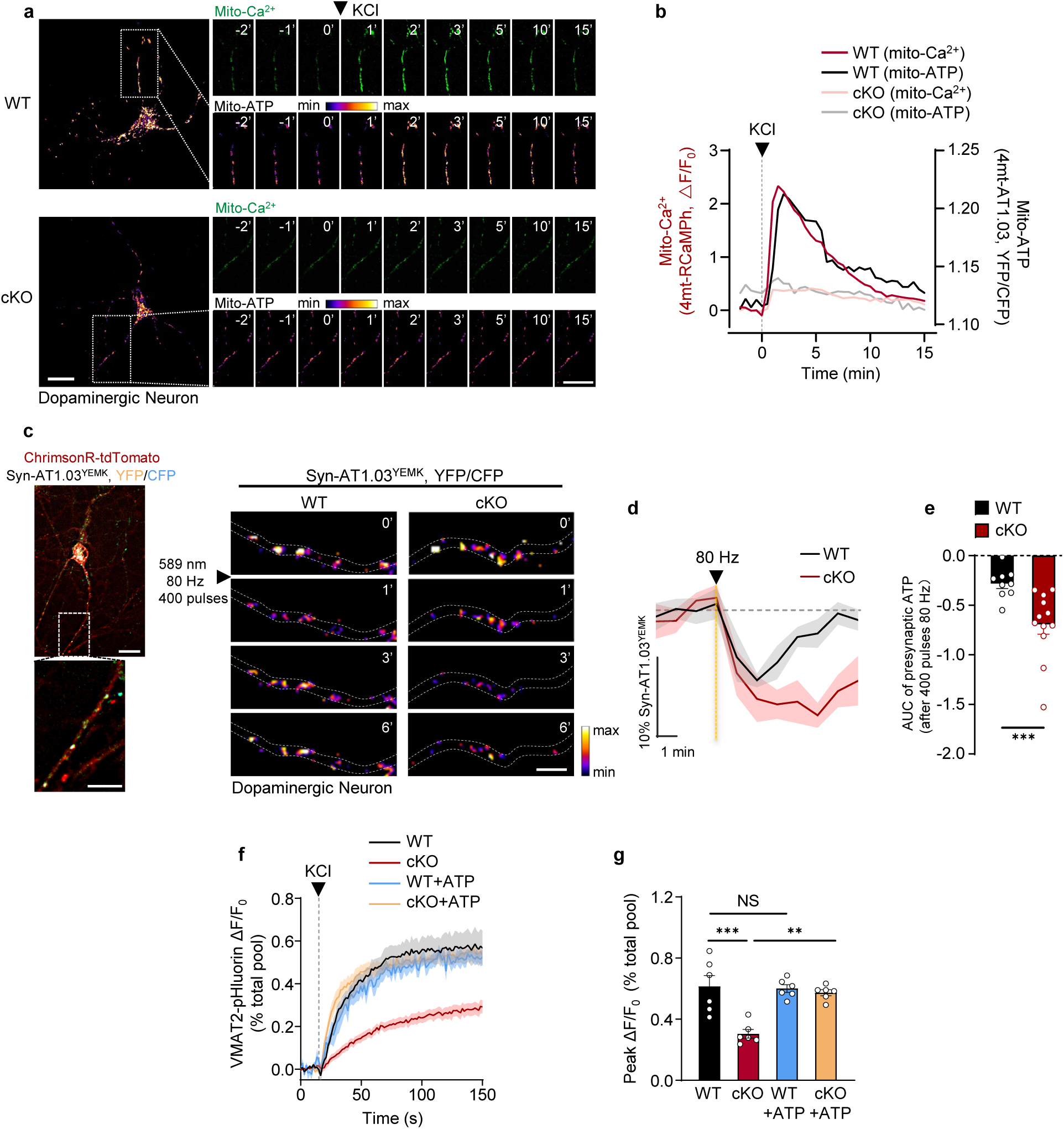
Mitochondrial calcium influx provides energy for sustained dopamine release. **a**, Pseudocolored time-lapse fluorescence images of a representative dopamine neuron co-expressing 4mt-RCaMPh (green) and 4mt-ATeam1.03 (pseudocolor) following KCl stimulation (75 mM). The color bar indicates the YFP/CFP emission ratio (pseudocolor). Scale bar: 10 μm. **b**, Representative traces of mitochondrial Ca^2+^ (4mt-RCaMPh) and mitochondrial ATP (4mt-ATeam1.03) dynamics, corresponding to the images in (**a**). **c**, (Left) Representative dopaminergic neuron co-expressing ChrimsonR-tdTomato and presynaptic ATP sensor Syn-AT1.03^YEMK^. The inset showing an axon expressing ATP probes at presynaptic sites. Scale bar: 10 μm (top), 5 μm (bottom). (Right) Representative pseudocolored time-lapse images showing presynaptic ATP dynamics (Syn-AT1.03^YEMK^) in dopaminergic axonal boutons from WT and DAT-*MCU* cKO neurons after 589 nm, 80 Hz, 400 pulses laser stimulation. Scale bar: 1 μm. Color bar indicates the YFP/CFP ratio. **d**, Normalized presynaptic ATP traces during 80 Hz laser stimulation (400 pulses) in WT (*n =* 9) and DAT-*MCU* cKO (*n =* 12) neurons. Grey dotted line: baseline; yellow dotted line: start of stimulation. Scale bar: 1 min (horizontal), 10% of ATP level change relative to baseline (vertical). **e**, Bar graph showing AUC of presynaptic ATP levels (Syn-AT1.03^YEMK^) during 7-min period after optogenetic stimulation (589 nm, 80 Hz, 400 pulses) in WT (*n =* 9) and DAT-*MCU* cKO (*n =* 12) neurons, based on the data in (**d**). **f**, Traces of VMAT2-pHluorin fluorescence in dopaminergic neurons following KCl stimulation, normalized to the total pool of dopamine exocytosis. (*n =* 6 cells per group). WT and DAT-*MCU* cKO neurons were pretreated with or without ATP (5 mM) for 60 min before imaging. **g**, Bar graphs showing normalized peak VMAT2-pHluorin fluorescence for dopamine exocytosis, based on the data in (**f**) (*n =* 6 cells per group). Data are presented as means ± SEM. Statistical analysis was performed using Mann-Whitney test (**e**) and one-way ANOVA with Bonferroni multiple comparison test (**g**). NS, not significant; \**P* < 0.05, \*\**P* < 0.01, \*\*\**P* < 0.001.

Given that dopamine release relies heavily on ATP-dependent vesicular transport and exocytosis at presynaptic terminals^20,25^, we next examined local ATP dynamics around synaptic vesicles. To assess local ATP fluctuations around vesicles during neurotransmission, we generated dopaminergic neurons expressing Synaptophysin-ATeam1.03^YEMK^ (Syn-AT1.03^YEMK^), a vesicle-localized ATP indicator^22,49^. Additionally, these neurons also expressed ChrimsonR, enabling optogenetic stimulation at various frequencies (5 Hz to 80 Hz) (Extended Data Fig. 3f,g). Time-lapse imaging revealed that high-frequency (80 Hz) optogenetic stimulation, which mimics intensive neuronal activation by drugs, led to a rapid decline of ATP levels at presynaptic vesicles. This decline was followed by prompt recovery within minutes in WT neurons. In contrast, MCU-deficient neurons exhibited a more pronounced ATP decline, and the recovery was significantly delayed (Fig. 4c–e). These results suggest a critical role of MCU-mediated rapid ATP production in compensating energy deficit caused by dopaminergic neuronal hyperactivation at presynapses.

To confirm that the reduced dopamine release observed in MCU-deficient neurons was due to insufficient ATP availability, we performed ATP supplementation experiments. We first confirmed that the free ATP levels in the axon of dopaminergic neurons were elevated following ATP supplementation (Extended Data Fig. 3h,i). Pretreating MCU-depleted dopaminergic neurons with ATP resulted in the restoration of impaired dopamine release (Fig. 4f,g). These results indicate that MCU-mediated mitochondrial Ca^2+^ influx regulates dopamine release by promoting mitochondrial ATP production.

Taken together, these findings suggest that drug-induced dopamine release is an energy-intensive process that relies on MCU-mediated Ca^2+^ entry into mitochondria to stimulate rapid ATP production. This newly synthesized ATP is crucial for replenishing the energy consumed at synaptic vesicles, thereby supporting the sustained dopamine release during intense neuronal activation.

### MCU-mediated mitochondrial Ca^2+^ influx selectively supports sustained high-intensity dopamine release

Our results suggest that mitochondrial Ca^2+^ influx is crucial for energy production necessary for high-intensity dopamine release. Notably, MCU deficiency did not alter basal ATP levels in dopaminergic terminals as indicated by the fluorescence of Syn-AT1.03^YEMK^ (Fig. 5a,b). High-performance liquid chromatography (HPLC) analysis of NAc tissues also showed no significant differences in basal dopamine levels between DAT-*MCU* cKO and WT mice (Fig. 5c-e and Extended Data Fig. 4a,b). These findings suggest that MCU might specifically regulate dopamine release under conditions of elevated energy demand, such as drug induced dopamine release, rather than maintaining basal dopamine transmission.

**Fig. 5.**
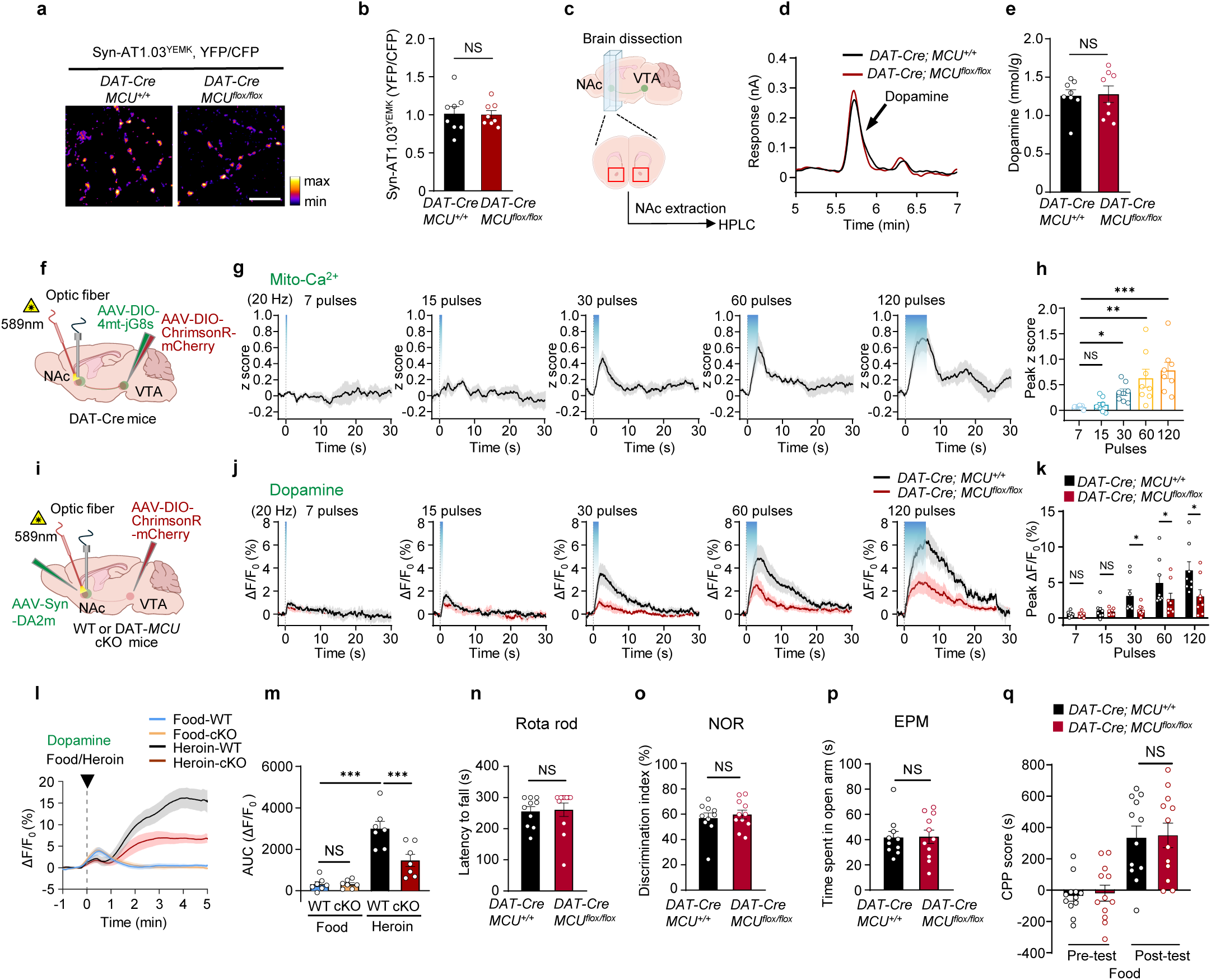
MCU knockout specifically blocks drug-induced excessive dopamine release without affecting dopamine-related physiological functions. **a,b**, Representative images (**a**) and quantification (**b**) showing that MCU deletion does not alter baseline ATP levels in dopaminergic neurons. Dopaminergic neurons from *DAT-Cre; MCU^+/+^* and *DAT-Cre; MCU^flox/flox^* mice were transfected with the presynaptic ATP probe Syn-AT1.03^YEMK^. The YFP/CFP ratio reflects presynaptic ATP level. Scale bar: 5 μm (*n =* 8 mice per group). **c**, Schematic showing dopamine content measurement in the NAc of mice using high-performance liquid chromatography (HPLC). **d**,**e**, Representative dopamine response curves (**d**) and average dopamine levels (**e**) in the NAc of WT and DAT-*MCU* cKO mice, measured by HPLC with electrochemical detection (*n =* 8 mice per group). **f**, Schematic showing fiber photometry setup for monitoring mitochondrial Ca^2^⁺ dynamics during optogenetic stimulation in DAT-Cre mice. **g**, Traces of mitochondrial Ca^2+^ dynamics (4mt-jGCaMP8, z score) in NAc dopaminergic terminals before and after 20 Hz optogenetic stimulation (blue shading) with 7, 15, 30, 60, or 120 pulses (*n* = 8 mice). **h**, Bar graph showing peak z score of the mitochondrial Ca^2+^ signals during optogenetic stimulation (20 Hz) with different pulse numbers, based on the data in (**g**) (*n* = 8 mice). **i**, Schematic showing fiber photometry setup for monitoring dopamine dynamics during optogenetic stimulation in WT and DAT-*MCU* cKO mice. **j**, Traces of dopamine signals (DA2m, ΔF/F_0_) in the NAc before and after 20 Hz optogenetic stimulation (blue shading) with 7, 15, 30, 60, or 120 pulses in WT and DAT-*MCU* cKO mice (*n* = 8 mice per group). **k**, Bar graph showing peak ΔF/F_0_ of the dopamine signals during optogenetic stimulation (20 Hz) with different pulse numbers in WT and DAT-*MCU* cKO mice, based on the data in (**j**) (*n* = 8 mice per group). **l**, Traces of dopamine signals (DA2m, ΔF/F_0_) in the NAc 1 min before and 5 min after food pellet intake (14 mg) or heroin injection (10 mg/kg) in WT and DAT-*MCU* cKO mice (*n =* 7 mice per group). Arrow indicates the start of food intake or injection. **m**, Bar graphs showing AUC of dopamine fluorescence signals (ΔF/F_0_) in the NAc during the 5-min period following food pellet intake (14 mg) or heroin injection in WT and DAT-*MCU* cKO mice (*n =* 7 mice per group), based on data from (**l**). **n–p**, Behavioral assessments show no differences between WT and DAT-MCU cKO mice in: latency to fall in the rota-rod test (**n**), discrimination index in the novel object recognition (NOR) task (**o**), time spent in the open arm of the elevated plus maze test (EPM) (**p**) (WT: *n* = 10, DAT-*MCU* cKO: *n* = 11 for all). **q**, CPP scores during pre-test and post-test sessions food in WT and DAT-*MCU* cKO mice (*n =* 12 per group). Data are presented as means ± SEM. Statistical analysis included two-sided unpaired Student’s t-test (**b**, **e**, **k**-[7, 15 pluses], **p**), Mann Whitney test (**k**-[30, 60, 120 pluses], **n**, **o**), Kruskal-Wallis test with a Dunn’s multiple comparison test (**h**), One-way ANOVA with Bonferroni multiple comparison test (**m**), and two-way ANOVA with Bonferroni post hoc test (**q**). NS, not significant; \**P* < 0.05, \*\**P* < 0.01, \*\*\**P* < 0.001.

VTA dopaminergic neurons exhibit two distinct firing patterns: low frequency tonic firing (1–8 Hz) and high-frequency phasic burst firing (>15 Hz)^42^. Tonic firing maintains baseline dopamine levels, while phasic firing mediates reward-related behaviors^50^. To examine whether MCU differentially regulates tonic and phasic firing-induced dopamine release, we utilized optogenetics combined with fiber photometry recordings. We expressed ChrimsonR in the VTA and DA2m in the NAc for optogenetic stimulation and dopamine detection, respectively, and co-expressed 4mt-jGCaMP8s with ChrimsonR in VTA dopaminergic neurons to measure terminal mitochondrial Ca^2+^ in the NAc. An optic fiber was implanted above the NAc for simultaneous stimulation and recordings (Extended Data Fig. 4c,d). We optogenetically stimulated dopaminergic terminals in the NAc with varying frequencies (4–40 Hz) using the same number of pulses (120 pulses) in both WT and DAT-*MCU* cKO mice. “Tonic” stimulation (4 Hz) only slightly increased dopamine release, with no significant difference between MCU-deficient and control mice. In contrast, “phasic” stimulation (20 Hz or 40 Hz) caused a marked increase in dopamine release in control mice, which was significantly reduced in MCU-deficient mice (Extended Data Fig. 4e–g). Consistent with the role of MCU exclusively in phasic dopamine release, mitochondrial Ca^2+^ levels were elevated during “phasic” stimulation, but not “tonic” stimulation (Extended Data Fig. 4h–j). These results demonstrate that MCU-mediated mitochondrial Ca^2+^ influx specifically occurs during phasic, but not tonic firing-induced dopamine release.

Having established MCU’s role in phasic firing-induced dopamine release, we next investigated whether different stimulation intensities affect mitochondrial Ca^2+^ dynamics by varying the number of pulses during 20 Hz optogenetic stimulation. Fewer pulses (7 and 15 pulses) did not significantly alter mitochondrial Ca^2+^ levels, while more pulses (30, 60, and 120 pulses) resulted in rapid and marked increases in mitochondrial Ca^2+^ levels (Fig. 5f–h). Consistently, optogenetic stimulation with fewer pulses (7 and 15 pulses at 20 Hz) elicited relatively low levels of dopamine release, with no significant differences between genotypes. However, more pulses (30, 60, and 120 pulses at 20 Hz) substantially increased dopamine release, and this response was significantly reduced in MCU-deficient mice (Fig. 5i–k). These findings demonstrate that MCU-mediated mitochondrial Ca^2+^ influx is specifically induced during high-intensity dopamine release, paralleling our *in vitro* observations that MCU is essential for intense dopamine release (Fig.4a–g).

### MCU knockout does not affect dopamine-related physiological functions in mice

Previous studies have shown that natural rewards, such as food intake, elicit significantly weaker dopamine release compared to drug stimulation^51^. To investigate the role of MCU in both natural and drug-induced reward behaviors, we monitored the dopamine levels in the NAc following food intake or heroin administration. While food intake resulted in a modest increase in dopamine levels, heroin administration led to a substantial dopamine surge (Fig. 5l,m). Notably, DAT-*MCU* cKO mice showed markedly reduced dopamine release in response to heroin, but not to food intake (Fig. 5l,m). Behavioral assessments revealed that these MCU-deficient mice exhibited normal performance in motor coordination, learning, memory, anxiety, and depression, as well as normal natural reward learning in the food-conditioned place preference (CPP) assay (Fig. 5n–q and Extended Data Fig. 5a–f). Collectively, these findings suggest that MCU-mediated mitochondrial Ca^2+^ influx is crucial specifically for drug-induced dopaminergic responses, while not affecting other dopamine-related behaviors.

To assess the broader impact of MCU deficiency, we generated pan-neuronal MCU knockout mice by crossing *MCU* floxed mice with *Nestin-Cre* mice (Extended Data Fig. 5g). These Nestin-*MCU* cKO mice were viable with normal Mendelian inheritance ratios (Extended Data Fig. 5h). When MCU was knocked out, the expression of TH and DAT, two markers of dopaminergic neurons, remained unaltered in midbrain and striatum (Extended Data Fig. 5i,j), indicating that MCU deficiency does not impair dopaminergic neurons. Moreover, Nestin-*MCU* cKO mice also exhibited normal motor coordination and learning functions (Extended Data Fig. 5k,l). These results collectively demonstrate that MCU knockout does not affect dopamine-related physiological functions in mice, suggesting that targeting MCU could be a promising and safe therapeutic strategy for drug addiction.

### Pharmacological inhibition of MCU alleviates drug reward behaviors in mice

Our recent study has identified berberine (BBR), a quaternary ammonia compound derived from plants, as a potent MCU inhibitor that specifically blocks mitochondrial Ca^2+^ uptake without affecting cytoplasmic Ca^2+^ levels in HeLa cells^52^. In cultured dopaminergic neurons, we confirmed that berberine dose-dependently inhibited KCl-induced mitochondrial Ca^2+^ elevation (Extended Data Fig. 6a,b), but did not affect cytosolic Ca^2+^ changes (Extended Data Fig. 6c). To further investigate the effects of berberine *in vivo*, we examined its impact on heroin-induced mitochondrial Ca^2+^ influx and dopamine release in the NAc. We used optic fibers with integrated microfluidic channels implanted above the NAc, allowing for simultaneous fluorescent signals recording and localized drug delivery (Fig. 6a). Intra-NAc administration of berberine significantly inhibited heroin-induced mitochondrial Ca^2+^ influx and dopamine release in the NAc (Fig. 6b,c). Moreover, although the baseline locomotion in mice remained unaffected (Extended Data Fig. 7a,b), berberine markedly reduced heroin-stimulated locomotor activity in wild-type mice (Fig. 6d,e). Notably, this effect was absent in DAT-*MCU* cKO mice (Fig. 6d,e), suggesting that berberine exerts its role via targeting MCU. Using the CPP paradigm (Extended Data Fig. 6d), we further demonstrated that intra-NAc berberine administration during the conditioning phase significantly attenuated the rewarding effects of heroin (Extended Data Fig. 6e), which was consistent with the phenotype observed in DAT-*MCU* cKO mice. These findings collectively indicate that berberine modulates drug reward by blocking MCU-mediated mitochondrial Ca^2+^ influx and subsequent dopamine release in the NAc.

**Fig. 6.**
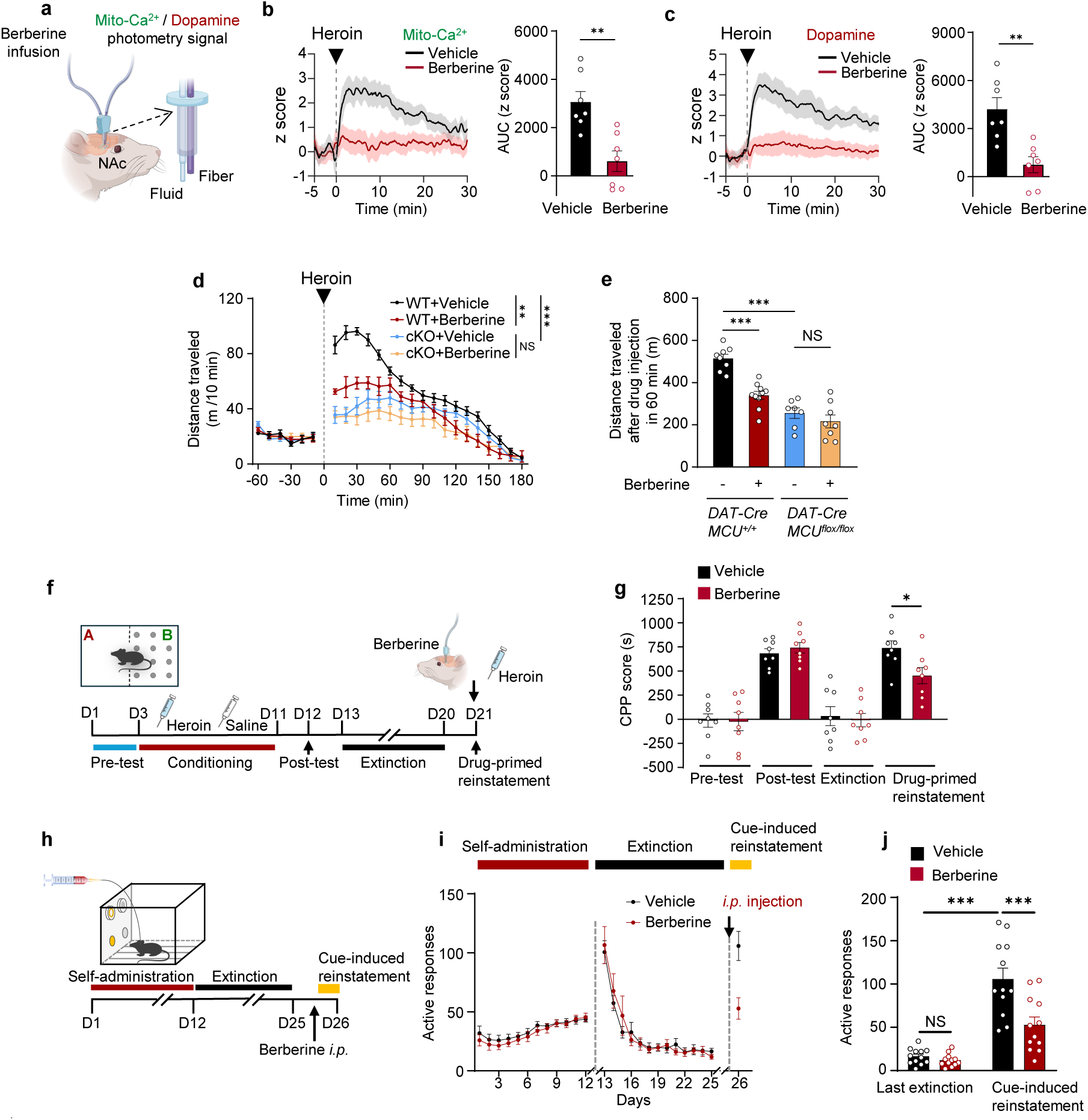
Pharmacological inhibition of MCU attenuates drug reward behaviors in mice. **a**, Schematic showing combined fiber photometry recording and local drug infusion setup using opto-fluid cannulas (OFC). **b**, Traces (left) and corresponding AUC analysis (30-min period, right) of mitochondrial Ca^2^⁺ dynamics (4mt-GCaMP8s fluorescence, z score) in NAc dopaminergic terminals aligned to heroin injection (grey dotted line and arrow) in vehicle- and berberine-pretreated (30 min) mice (*n =* 8 per group). **c**, Traces (left) and corresponding AUC analysis (30-min period, right) of dopamine signals (rDA3m, z score) in the NAc aligned to heroin injection (grey dotted line and arrow) in vehicle- and berberine-pretreated (30 min) mice (*n =* 8 per group). **d**,**e**, Distance traveled in open field before and after heroin injection (10 mg/kg, grey dotted line) in *DAT-Cre; MCU^+/+^* (WT) mice with intra-NAc vehicle (black, *n =* 8) or berberine (red, *n =* 10), and *DAT-Cre; MCU^flox/flox^* (cKO) mice with vehicle (blue, *n* = 7) or berberine (yellow, *n =* 8), shown as time course (**d**) and cumulative distance during 60-min post-injection period (**e**). **f**, Schematic of CPP paradigm showing conditioning, extinction, and reinstatement phases. Berberine (0.3 μg/side) was infused into NAc 30 min before reinstatement test. **g**, CPP scores during pre-test, post-test, extinction, and drug-primed reinstatement in mice receiving intra-NAc vehicle (black) or berberine (red) before reinstatement test (n = 8 per group). **h**, Schematic of cocaine self-administration paradigm showing cocaine self-administration training, extinction, and cue-induced seeking test. Berberine (3 mg/kg, i.p.) was systemically administered 120 min prior to the cue-induced reinstatement test session. **i**, Active nose-poke responses during cocaine self-administration training, extinction, and cue-induced reinstatement test in vehicle- and berberine-treated mice (n = 12 per group). Arrows indicate intraperitoneal administration of vehicle or berberine 120 min prior to the cue-induced reinstatement test. **j**, Active nose-poke responses between vehicle and berberine groups during the final extinction session and cue-induced reinstatement test of cocaine self-administration. Data are presented as means ± SEM. Statistical analysis included two-sided unpaired Student’s t-test (**b**, **c**), one-way ANOVA with Bonferroni multiple comparison test (**e**), and two-way ANOVA with Bonferroni post hoc test (**d**, **g, i, j**). NS, not significant; \**P* < 0.05, \*\**P* < 0.01, \*\*\**P* < 0.001.

The most challenging aspect of substance use disorders is the high propensity for relapse, such as drug-induced, cue-induced, and context-induced relapse, during which dopamine release in the NAc is markedly elevated^53–55^. To evaluate whether targeting MCU-mediated pathological dopamine release could potentially prevent relapse, we employed a confined CPP reinstatement paradigm to investigate whether inhibition of MCU by berberine treatment affects drug-induced reinstatement (Fig. 6f). A single intra-NAc injection of berberine before the CPP test significantly inhibited the reinstatement of heroin-CPP (Fig. 6g). Beyond drug-primed relapse, cue-induced craving represents another major trigger for relapse. We next used the cocaine self-administration paradigm, a well-established model for assessing cue-induced drug-seeking behavior (Fig. 6h), where cue exposure triggers phasic dopamine release in the NAc, leading to relapse^53,56,57^. Notably, this relapse is driven by cue-induced dopamine release in the NAc, irrespective of the drug type. Following stable cocaine self-administration training (Fig. 6i), mice underwent a two-week extinction period before the reinstatement test. Mice were then randomly assigned to either the vehicle group or the berberine-treated group. Vehicle-treated mice showed a significant increase in active nose-pokes during the cue presentation compared to their extinction baseline, validating the reinstatement model. Importantly, systemic administration of berberine (3 mg/kg, i.p.) 2h before the test significantly attenuated this cue-induced drug-seeking behavior (Fig. 6j). To confirm the specificity of berberine’s effect on cue-induced seeking behavior given its systemic administration, we employed an inducible conditional knockout strategy to selectively delete MCU in dopaminergic neurons prior to the drug-seeking test (Extended Data Fig. 6f,g). Consistent with the pharmacological intervention results, genetic ablation of MCU specifically in dopaminergic neurons significantly suppressed cue-induced drug-seeking behavior (Extended Data Fig. 6h,i). In addition to efficacy, we evaluated the safety profile of berberine. Intra-NAc injection of berberine did not cause anxiety, depression, or cognitive deficits (Extended Data Fig. 7a,c–h), consistent with the observations in DAT-*MCU* cKO and Nestin-*MCU* cKO mice. Collectively, these data suggest that berberine is a safe and effective pharmacological agent for reducing heroin-CPP acquisition and preventing drug-induced relapse. This highlights berberine’s potential as a novel therapeutic strategy for treating drug addiction without adversely affecting other physiological functions.

## Discussion

By visualizing mitochondrial Ca^2+^ influx in a drug- and brain region-specific manner following drug administration in mice, we have uncovered a critical mechanism through which drugs of abuse such as opioids hijack the body’s energy metabolism to drive drug reward and addiction. Our findings demonstrate that mitochondrial Ca^2+^ influx plays a critical role in meeting the elevated energy demands required for intense dopamine release during drug-induced reward and addiction. This highlights a previously underappreciated link between energy metabolism and addiction mechanisms. These insights provide a novel perspective on addiction biology and suggest that targeting mitochondrial Ca^2+^ influx and the associated bioenergetic processes may represent a promising and safer therapeutic strategy for treating drug addiction.

The brain, particularly regions with high synaptic activity, is exceptionally energy-demanding, with synaptic transmission and ion gradient maintenance consuming the majority of neural energy^20,58–60^. Recent evidence hints at the involvement of energy-related metabolic processes in addiction regulation. For instance, chronic exposure to addictive drugs has been shown to trigger mitochondrial fragmentation in neurons, resulting in impaired mitochondrial function^61,62^. Additionally, astrocyte-neuron metabolic coupling through lactate shuttle within the amygdala has been implicated in the reconsolidation of drug-associated memories and the persistence of craving^63,64^. In this study, we provide direct evidence that drug-induced mitochondrial ATP production is indispensable for sustaining the phasic dopamine release necessary for drug reward and addiction. Hyperactivation of mesolimbic circuits induces sustained dopamine release that imposes significant bioenergetic demands and causes severe energy deficit in dopaminergic terminals. In response, mitochondria within the NAc dopaminergic terminals initiate sustained Ca^2+^ influx, rapidly activating an emergency energy supply mechanism to maintain this pathological neurotransmitter release. These findings support a model wherein addictive drugs actively reprogram mitochondrial bioenergetics, specifically to meet the extreme demands of addiction-induced neurotransmitter surges. By uncovering this direct link between cellular energetics and the core mechanisms of addiction, our research introduces a fundamentally new perspective on the neurobiology of addiction. While our optogenetic approaches provided valuable insights into activity-dependent energy dynamics in dopaminergic neurons in vitro, they cannot fully replicate the complex in vivo conditions involving drug metabolism and circuit-level interactions. Future studies employing more sensitive ATP biosensors should monitor spatiotemporal ATP dynamics across different neuronal populations during addiction, providing a more comprehensive understanding of bioenergetic regulation in addiction pathogenesis.

Calcium signaling plays a critical role in regulating neuronal activity^30,31,65^. Previous studies indicate that certain drugs activate dopaminergic neurons^26,42^, increasing Ca^2+^ influx to stimulate dopamine release and synaptic plasticity^66–68^— foundational mechanisms for reinforcing addictive behaviors and consolidating addiction memories^26,69^. Although cytosolic Ca^2+^ elevation typically triggers mitochondrial Ca^2+^ uptake^43,44^, the potential involvement of mitochondrial Ca^2+^ influx in addiction has remained unexplored. Our findings reveal that heroin elevates cytosolic Ca^2+^ in both the VTA soma and NAc terminals, yet mitochondrial Ca^2+^ influx occurs exclusively in NAc terminals, likely due to the high mitochondrial density in this region^24^. This region-specific Ca^2+^ influx enables the release of large amounts of dopamine, thereby reinforcing drug reward. This finding extends the role of Ca^2+^ signaling in addiction regulation from the cytosol to mitochondria. Drug-induced mitochondrial Ca^2+^ influx at dopaminergic synaptic terminals enhances neurotransmitter release through energy metabolism support, consistent with previous *in vitro* studies of high-frequency electrical stimulation^20,70^. This process is unlikely to simply buffer cytoplasmic Ca^2+^, as cytoplasmic Ca^2+^ buffering can hinder neuronal activation^71,72^. Although our study highlights the unique function of drug-induced mitochondrial Ca^2+^ influx in dopaminergic neurons, future studies should investigate the generality and spatiotemporal characteristics of this mechanism in other addiction-related cell types (e.g., glutamatergic neurons, GABAergic neurons, glial cells) and brain regions (e.g., hippocampus, amygdala). This exploration should also include different stages of addiction, such as withdrawal and relapse, to fully elucidate the role of mitochondrial Ca^2+^ influx throughout addiction cycle.

Substance addiction remains a major public health issue^11^, and current treatment strategies face significant challenges due to the lack of ideal pharmacological targets and the complexity of addiction mechanisms^73–75^. Reducing drug reward effects and inhibiting drug-seeking behaviors, both of which critically depend on the activation of dopamine neurons, are key goals in addiction treatment^76–78^. However, existing dopamine-modulating therapies in clinical and preclinical setting, such as opioid receptor or dopamine receptor inhibition, often lead to broad disruption of reward circuitry and various side effects, including anhedonia and motivational deficits^16,79–81^. Our study identifies a novel mechanism where MCU-mediated mitochondrial Ca^2+^ influx selectively regulates drug-induced dopamine release, a critical process in drug reward. Importantly, this mechanism is specifically activated during drug use but remains inactive during natural rewards, providing a unique insight into the selective nature of addiction-related dopamine release. Moreover, while we did not provide direct evidence that drug-related cues induce mitochondrial Ca^2+^ influx in the NAc, our self-administration experiments demonstrate that MCU knockout or inhibition significantly reduces cue-trigger relapse behaviors. This finding, along with previous reports that cues trigger phasic dopamine release to facilitate relapse by enhancing drug-related memories and motivational drives^39,42^, highlights MCU’s pivotal role in addiction-related processes. Unlike current strategies, MCU modulation selectively disrupts relapse mechanisms without the side effects typically associated with broad dopamine system manipulation. This specificity is further supported by the fact that MCU is primarily involved in responding to acute energy demands, such as during exercise or cell division^36,82^, rather than playing a role in basal mitochondrial respiration^70,82,83^. Therefore, MCU targeting provides a promising and innovative strategy to manage drug reward and reduce drug-seeking behaviors, with the potential for fewer side effects.

Our study reveals the therapeutic potential of MCU inhibition for treating addiction. Through experiments with berberine, a natural compound known for its safety profile^84^, we demonstrate that it effectively suppresses excessive dopamine release via MCU inhibition in the NAc and reduces cue-triggered relapse. This provides a molecular basis for previously reported effects of berberine on addictive behaviors^85,86^ and validates MCU as a promising target for addiction treatment. Notably, not all drugs in our study induced mitochondrial Ca²^+^ influx. Only substances that promote dopamine release, such as opioids, exhibited this effect. Methamphetamine, primarily inhibiting dopamine reuptake^10^, also promotes dopamine release to a lesser extent^48^, exhibiting in weaker mitochondrial Ca²^+^ influx. In contrast, cocaine, predominantly acting by inhibiting dopamine reuptake^10^, does not induce mitochondrial Ca^2+^ influx. These findings underscore the specific role of dopamine release-promoting substances in driving mitochondrial Ca²^+^ influx and suggest that MCU inhibition could be particularly effective for treating addiction to these substances. Consequently, MCU-targeted strategies could attenuate drug reward induced by heroin, morphine, methamphetamine, as well as addiction to other dopamine release-promoting substances like alcohol^87^, nicotine^88^, and fentanyl^89^. Further investigation into MCU inhibition in preclinical models and clinical applications is warranted.

In summary, our study reveals a previously unrecognized mechanism in which mitochondrial Ca^2^LJ influx through the MCU actively fuels the excessive dopamine surges that drive addiction. By identifying MCU as a key regulator of addiction-specific energy dynamics, our work opens new therapeutic avenues with the potential to disrupt the metabolic basis of drug reward while preserving natural reward processes. Furthermore, this finding redefines energy metabolism in the brain, not merely as a background process supporting physiological brain functions, but as an active and essential driver of pathological processes such as addiction. This encourages further exploration of diverse energy metabolism mechanisms that may underlie other neurological and psychiatric disorders, expanding our understanding of disease pathology and fostering the development of novel therapeutic approaches.

## Acknowledgments

We thank Jin Li and Ning Wu for providing reagents; Cailing Wang for assistance with experiments. We thank Hongtao Yu for helpful discussion. We thank Kai Wang and Xin Xu for assistance with microscopy assays. This work was funded by the National Natural Science Foundation of China (82025028, 92354304, and 82000107) and the National Key Research and Development Program of China (2021YFA1300203 and 2020YFA0113300).

## Author Contributions

X.P., Teng L., A.L.L., and J.G. contributed to study design. Teng L., J.G., X.H., and L.Z. performed the photometry and two-photon data acquisition and analyses. J.G., L.Z., W.W., and X.W. contributed to viral injections and behavioral experiments. H.Z., L.Z., and Teng L. performed neuron culture experiments. G.L. and X.H. performed biochemistry and immunostaining experiments. J.C., Ting L., C.L., S.C., J.Y., T.Z., A.L.L., and X.M.Z. contributed to interpreting the results. X.P., Teng L., and A.L.L. supervised the research. X.P. wrote the manuscript with the assistance of Teng L., J.G., H.Z., and X.H.

## Declaration of Interests

The authors declare no competing interests.

## Notes

### Competing Interest Statement

The authors have declared no competing interest.

## References

1 Stuber, G. D. Neurocircuits for motivation. Science 382, 394–398 (2023).

2 Liu, W. W. & Bohorquez, D. V. The neural basis of sugar preference. Nat Rev Neurosci 23, 584–595 (2022).

3 Sosa, M. & Giocomo, L. M. Navigating for reward. Nat Rev Neurosci 22, 472–487 (2021).

4 Dai, B. et al. Responses and functions of dopamine in nucleus accumbens core during social behaviors. Cell Rep 40, 111246 (2022).

5 Mohebi, A. et al. Dissociable dopamine dynamics for learning and motivation. Nature 570, 65–70 (2019).

6 Berridge, K. C. & Kringelbach, M. L. Pleasure systems in the brain. Neuron 86, 646–664 (2015).

7 Tan, B. et al. Drugs of abuse hijack a mesolimbic pathway that processes homeostatic need. Science 384, eadk6742 (2024).

8 Koob, G. F. & Volkow, N. D. Neurobiology of addiction: a neurocircuitry analysis. Lancet Psychiatry 3, 760–773 (2016).

9 Volkow, N. D. & Morales, M. The Brain on Drugs: From Reward to Addiction. Cell 162, 712–725 (2015).

10 Covey, D. P., Roitman, M. F. & Garris, P. A. Illicit dopamine transients: reconciling actions of abused drugs. Trends Neurosci 37, 200–210 (2014).

11 Volkow, N. D. & Blanco, C. Substance use disorders: a comprehensive update of classification, epidemiology, neurobiology, clinical aspects, treatment and prevention. World Psychiatry 22, 203–229 (2023).

12 Volkow, N. D. & Boyle, M. Neuroscience of Addiction: Relevance to Prevention and Treatment. Am J Psychiatry 175, 729–740 (2018).

13 Ewing, S. T. et al. Low-dose polypharmacology targeting dopamine D1 and D3 receptors reduces cue-induced relapse to heroin seeking in rats. Addict Biol 26, e12988 (2021).

14 Caine, S. B. et al. Lack of self-administration of cocaine in dopamine D1 receptor knock-out mice. J Neurosci 27, 13140–13150 (2007).

15 Wang, J. et al. Microglial activation contributes to depressive-like behavior in dopamine D3 receptor knockout mice. Brain Behav Immun 83, 226–238 (2020).

16 Nutt, D. J., Lingford-Hughes, A., Erritzoe, D. & Stokes, P. R. The dopamine theory of addiction: 40 years of highs and lows. Nat Rev Neurosci 16, 305–312 (2015).

17 Leucht, S. et al. Comparative efficacy and tolerability of 15 antipsychotic drugs in schizophrenia: a multiple-treatments meta-analysis. Lancet 382, 951–962 (2013).

18 Dienel, G. A. Brain Glucose Metabolism: Integration of Energetics with Function. Physiol Rev 99, 949–1045 (2019).

19 Harris, J. J., Jolivet, R. & Attwell, D. Synaptic energy use and supply. Neuron 75, 762–777 (2012).

20 Li, S. & Sheng, Z. H. Energy matters: presynaptic metabolism and the maintenance of synaptic transmission. Nat Rev Neurosci 23, 4–22 (2022).

21 Sudhof, T. C. Neurotransmitter release: the last millisecond in the life of a synaptic vesicle. Neuron 80, 675–690 (2013).

22 Pathak, D. et al. The role of mitochondrially derived ATP in synaptic vesicle recycling. J Biol Chem 290, 22325–22336 (2015).

23 Rangaraju, V., Calloway, N. & Ryan, T. A. Activity-driven local ATP synthesis is required for synaptic function. Cell 156, 825–835 (2014).

24 Sheng, Z. H. & Cai, Q. Mitochondrial transport in neurons: impact on synaptic homeostasis and neurodegeneration. Nat Rev Neurosci 13, 77–93 (2012).

25 Devine, M. J. & Kittler, J. T. Mitochondria at the neuronal presynapse in health and disease. Nat Rev Neurosci 19, 63–80 (2018).

26 Luscher, C. & Malenka, R. C. Drug-evoked synaptic plasticity in addiction: from molecular changes to circuit remodeling. Neuron 69, 650–663 (2011).

27 Parvaz, M. A., Alia-Klein, N., Woicik, P. A., Volkow, N. D. & Goldstein, R. Z. Neuroimaging for drug addiction and related behaviors. Rev Neurosci 22, 609–624 (2011).

28 Moningka, H. et al. Can neuroimaging help combat the opioid epidemic? A systematic review of clinical and pharmacological challenge fMRI studies with recommendations for future research. Neuropsychopharmacology 44, 259–273 (2019).

29 Volkow, N. D., Fowler, J. S. & Wang, G. J. The addicted human brain: insights from imaging studies. J Clin Invest 111, 1444–1451 (2003).

30 Grienberger, C. & Konnerth, A. Imaging calcium in neurons. Neuron 73, 862–885 (2012).

31 Ghosh, A. & Greenberg, M. E. Calcium signaling in neurons: molecular mechanisms and cellular consequences. Science 268, 239–247 (1995).

32 Garbincius, J. F. & Elrod, J. W. Mitochondrial calcium exchange in physiology and disease. Physiol Rev 102, 893–992 (2022).

33 Rizzuto, R., De Stefani, D., Raffaello, A. & Mammucari, C. Mitochondria as sensors and regulators of calcium signalling. Nat Rev Mol Cell Biol 13, 566–578 (2012).

34 Territo, P. R., Mootha, V. K., French, S. A. & Balaban, R. S. Ca(2+) activation of heart mitochondrial oxidative phosphorylation: role of the F(0)/F(1)-ATPase. Am J Physiol Cell Physiol 278, C423–435 (2000).

35 McCormack, J. G., Halestrap, A. P. & Denton, R. M. Role of calcium ions in regulation of mammalian intramitochondrial metabolism. Physiol Rev 70, 391–425 (1990).

36 Zhao, H. et al. AMPK-mediated activation of MCU stimulates mitochondrial Ca(2+) entry to promote mitotic progression. Nat Cell Biol 21, 476–486 (2019).

37 Tarasova, N. V., Vishnyakova, P. A., Logashina, Y. A. & Elchaninov, A. V. Mitochondrial Calcium Uniporter Structure and Function in Different Types of Muscle Tissues in Health and Disease. Int J Mol Sci 20 (2019).

38 Bromberg-Martin, E. S., Matsumoto, M. & Hikosaka, O. Dopamine in motivational control: rewarding, aversive, and alerting. Neuron 68, 815–834 (2010).

39 Keiflin, R. & Janak, P. H. Dopamine Prediction Errors in Reward Learning and Addiction: From Theory to Neural Circuitry. Neuron 88, 247–263 (2015).

40 Coddington, L. T., Lindo, S. E. & Dudman, J. T. Mesolimbic dopamine adapts the rate of learning from action. Nature 614, 294–302 (2023).

41 Nestler, E. J. Is there a common molecular pathway for addiction? Nat Neurosci 8, 1445–1449 (2005).

42 Volkow, N. D., Wise, R. A. & Baler, R. The dopamine motive system: implications for drug and food addiction. Nat Rev Neurosci 18, 741–752 (2017).

43 Williams, G. S., Boyman, L., Chikando, A. C., Khairallah, R. J. & Lederer, W. J. Mitochondrial calcium uptake. Proc Natl Acad Sci U S A 110, 10479–10486 (2013).

44 Foskett, J. K. & Philipson, B. The mitochondrial Ca(2+) uniporter complex. J Mol Cell Cardiol 78, 3–8 (2015).

45 De Stefani, D., Raffaello, A., Teardo, E., Szabo, I. & Rizzuto, R. A forty-kilodalton protein of the inner membrane is the mitochondrial calcium uniporter. Nature 476, 336–340 (2011).

46 Baughman, J. M. et al. Integrative genomics identifies MCU as an essential component of the mitochondrial calcium uniporter. Nature 476, 341–345 (2011).

47 Sun, F. et al. Next-generation GRAB sensors for monitoring dopaminergic activity in vivo. Nat Methods 17, 1156–1166 (2020).

48 Hedges, D. M. et al. Methamphetamine Induces Dopamine Release in the Nucleus Accumbens Through a Sigma Receptor-Mediated Pathway. Neuropsychopharmacology 43, 1405–1414 (2018).

49 Li, S., Xiong, G. J., Huang, N. & Sheng, Z. H. The cross-talk of energy sensing and mitochondrial anchoring sustains synaptic efficacy by maintaining presynaptic metabolism. Nat Metab 2, 1077–1095 (2020).

50 Schultz, W. Behavioral dopamine signals. Trends Neurosci 30, 203–210 (2007).

51 Koob, G. F. & Volkow, N. D. Neurocircuitry of addiction. Neuropsychopharmacology 35, 217–238 (2010).

52 Zhao, H. et al. Berberine is a novel mitochondrial calcium uniporter (MCU) inhibitor that disrupts MCU-EMRE assembly. bioRxiv, 2024.2008.2018.607892 (2024).

53 Lujan, M. A. et al. A multivariate regressor of patterned dopamine release predicts relapse to cocaine. Cell Rep 42, 112553 (2023).

54 Liu, Y. & McNally, G. P. Dopamine and relapse to drug seeking. J Neurochem 157, 1572–1584 (2021).

55 Madayag, A. et al. Drug-induced plasticity contributing to heightened relapse susceptibility: neurochemical changes and augmented reinstatement in high-intake rats. J Neurosci 30, 210–217 (2010).

56 Jing, M. Y. et al. Activation of mesocorticolimbic dopamine projections initiates cue-induced reinstatement of reward seeking in mice. Acta Pharmacol Sin 43, 2276–2288 (2022).

57 Saunders, B. T., Yager, L. M. & Robinson, T. E. Cue-evoked cocaine “craving”: role of dopamine in the accumbens core. J Neurosci 33, 13989–14000 (2013).

58 Raichle, M. E. & Gusnard, D. A. Appraising the brain’s energy budget. Proc Natl Acad Sci U S A 99, 10237–10239 (2002).

59 Watts, M. E., Pocock, R. & Claudianos, C. Brain Energy and Oxygen Metabolism: Emerging Role in Normal Function and Disease. Front Mol Neurosci 11, 216 (2018).

60 Faria-Pereira, A. & Morais, V. A. Synapses: The Brain’s Energy-Demanding Sites. Int J Mol Sci 23 (2022).

61 Chandra, R. et al. Drp1 Mitochondrial Fission in D1 Neurons Mediates Behavioral and Cellular Plasticity during Early Cocaine Abstinence. Neuron 96, 1327–1341 e1326 (2017).

62 Jiang, C. et al. Targeting mitochondrial dynamics of morphine-responsive dopaminergic neurons ameliorates opiate withdrawal. J Clin Invest 134 (2024).

63 Boury-Jamot, B. et al. Disrupting astrocyte-neuron lactate transfer persistently reduces conditioned responses to cocaine. Mol Psychiatry 21, 1070–1076 (2016).

64 Chen, W. et al. Disrupting astrocyte-neuron lactate transport prevents cocaine seeking after prolonged withdrawal. Sci Adv 9, eadi4462 (2023).

65 Kawamoto, E. M., Vivar, C. & Camandola, S. Physiology and pathology of calcium signaling in the brain. Front Pharmacol 3, 61 (2012).

66 Inglebert, Y., Aljadeff, J., Brunel, N. & Debanne, D. Synaptic plasticity rules with physiological calcium levels. Proc Natl Acad Sci U S A 117, 33639–33648 (2020).

67 Neher, E. & Sakaba, T. Multiple roles of calcium ions in the regulation of neurotransmitter release. Neuron 59, 861–872 (2008).

68 Sudhof, T. C. Calcium control of neurotransmitter release. Cold Spring Harb Perspect Biol 4, a011353 (2012).

69 Kauer, J. A. & Malenka, R. C. Synaptic plasticity and addiction. Nat Rev Neurosci 8, 844–858 (2007).

70 Datta, S. & Jaiswal, M. Mitochondrial calcium at the synapse. Mitochondrion 59, 135–153 (2021).

71 Kwon, S. K. et al. LKB1 Regulates Mitochondria-Dependent Presynaptic Calcium Clearance and Neurotransmitter Release Properties at Excitatory Synapses along Cortical Axons. PLoS Biol 14, e1002516 (2016).

72 Vaccaro, V., Devine, M. J., Higgs, N. F. & Kittler, J. T. Miro1-dependent mitochondrial positioning drives the rescaling of presynaptic Ca2+ signals during homeostatic plasticity. EMBO Rep 18, 231–240 (2017).

73 Mehta, D. D. et al. A systematic review and meta-analysis of neuromodulation therapies for substance use disorders. Neuropsychopharmacology 49, 649–680 (2024).

74 Swinford-Jackson, S. E. et al. The Persistent Challenge of Developing Addiction Pharmacotherapies. Cold Spring Harb Perspect Med 11 (2021).

75 Oliva, E. M., Maisel, N. C., Gordon, A. J. & Harris, A. H. Barriers to use of pharmacotherapy for addiction disorders and how to overcome them. Curr Psychiatry Rep 13, 374–381 (2011).

76 Volkow, N. D., Michaelides, M. & Baler, R. The Neuroscience of Drug Reward and Addiction. Physiol Rev 99, 2115–2140 (2019).

77 Volkow, N. D., Fowler, J. S., Wang, G. J., Swanson, J. M. & Telang, F. Dopamine in drug abuse and addiction: results of imaging studies and treatment implications. Arch Neurol 64, 1575–1579 (2007).

78 Liu, X. et al. Preventing incubation of drug craving to treat drug relapse: from bench to bedside. Mol Psychiatry 28, 1415–1429 (2023).

79 Lutz, P. E. & Kieffer, B. L. Opioid receptors: distinct roles in mood disorders. Trends Neurosci 36, 195–206 (2013).

80 Salamone, J. D. & Correa, M. The mysterious motivational functions of mesolimbic dopamine. Neuron 76, 470–485 (2012).

81 Wise, R. A. Dopamine, learning and motivation. Nat Rev Neurosci 5, 483–494 (2004).

82 Pan, X. et al. The physiological role of mitochondrial calcium revealed by mice lacking the mitochondrial calcium uniporter. Nat Cell Biol 15, 1464–1472 (2013).

83 Alevriadou, B. R. et al. Molecular nature and physiological role of the mitochondrial calcium uniporter channel. Am J Physiol Cell Physiol 320, C465–C482 (2021).

84 Ju, J., Li, J., Lin, Q. & Xu, H. Efficacy and safety of berberine for dyslipidaemias: A systematic review and meta-analysis of randomized clinical trials. Phytomedicine 50, 25–34 (2018).

85 Shen, X. et al. Berberine Facilitates Extinction of Drug-Associated Behavior and Inhibits Reinstatement of Drug Seeking. Front Pharmacol 11, 476 (2020).

86 Bhutada, P. et al. Inhibitory effect of berberine on the motivational effects of ethanol in mice. Prog Neuropsychopharmacol Biol Psychiatry 34, 1472–1479 (2010).

87 Egervari, G., Siciliano, C. A., Whiteley, E. L. & Ron, D. Alcohol and the brain: from genes to circuits. Trends Neurosci 44, 1004–1015 (2021).

88 Pidoplichko, V. I., DeBiasi, M., Williams, J. T. & Dani, J. A. Nicotine activates and desensitizes midbrain dopamine neurons. Nature 390, 401–404 (1997).

89 Heilig, M. & Petrella, M. Neural pathways for reward and relief promote fentanyl addiction. Nature 630, 38–39 (2024).

