## Supplementary material for "Mitochondrial calcium influx-driven bioenergetics in the dopaminergic system selectively enable drug reward": Methods

### Animals

All the mouse lines used in this work were backcrossed and maintained on a C57BL/6N background. The generation of Mitochondrial Calcium Uniporter (MCU) conditional knockout mice (*MCU*$^{flox/flox}$) was achieved through targeted homologous recombination in embryonic stem (ES) cells. Specifically, loxP sites were integrated around exon 5 of the MCU gene, in collaboration with Cyagen Biosciences. To selectively delete the MCU in dopaminergic neurons, we initially crossed *DAT-IRES-Cre* (Jax 006660) mice with *MCU*$^{flox/flox}$ mice to generate *DAT-Cre*$^{+/-}$*; MCU*$^{flox/+}$ and *DAT-Cre*$^{-/-}$*; MCU*$^{flox/+}$ offspring. Subsequently, *DAT-Cre*$^{+/-}$*; MCU*$^{flox/flox}$ mice were generated by intercrossing *DAT-Cre*$^{+/-}$*; MCU*$^{flox/+}$ with *DAT-Cre*$^{-/-}$*; MCU*$^{flox/+}$ mice. Littermate *DAT-Cre*$^{+/-}$*; MCU*$^{+/+}$ mice served as controls. For global brain MCU deletion, *Nes-Cre*$^{+/-}$*; MCU*$^{flox/flox}$ mice were generated by crossing *Nestin-Cre* (Jax 003771) mice with *MCU*$^{flox/flox}$ mice for at least two generations, and the littermate *Nes-Cre*$^{-/-}$*; MCU*$^{flox/flox}$ mice served as controls. To restrict MCU deletion to dopamine neurons in the VTA, AAV2/5-mTH-NLS-Cre virus was injected into the VTA of *MCU*$^{flox/flox}$ mice. Adult C57BL/6N mice were obtained from Beijing Vital River Laboratory Animal Technology Co., Ltd. All fiber photometry and behavioral studies were conducted using adult mice (aged 8–12 weeks), with a balanced representation of both sexes in the cohort. Animals were housed in a specific-pathogen-free environment under a 12-hour light/dark cycle (50 lux, lights on 7:00 to 19:00), with regulated temperature (22°C) and humidity (40–60%). Food and water were available ad libitum, unless otherwise noted. All animal procedures were in accordance with the Institutional Animal Care and Use Committee of the National Center of Biomedical Analysis.

### Drugs

Morphine hydrochloride and cocaine hydrochloride were purchased from Qinghai Pharmaceutical Co., Ltd. (Xining, China). Heroin and methamphetamine (METH) were kindly provided by Dr. Jin Li and Dr. Ning Wu from the Beijing Institute of Pharmacology and Toxicology (Beijing, China). All addictive substances were dissolved in sterile 0.9% (w/v) sodium chloride solution immediately prior to use. Berberine (Shanghai Macklin Biochemical Co., Ltd., Shanghai, China) was dissolved in ultra-pure water, and heparin (Shanghai Macklin Biochemical Co., Ltd., Shanghai, China) was dissolved in 0.9% (w/v) sodium chloride solution. Both solutions were sterilized by filtration through a 0.22 μm pore size membrane filter before use.

**Plasmid and virus generation**

In experiments involving the stereotaxic injection of AAV into mice, the following viral vectors were used: AAV2/9-EF1a-DIO-4mt-jGCaMP8s ($3.3\times10^{12}$ genome copies/ml) from Obio Technology, Shanghai, China; AAV2/9-EF1a-DIO-GCaMP6m ($5.18\times10^{12}$ genome copies/ml), AAV2/5-mTH-NLS-Cre (AAV-TH-Cre, $5.88\times10^{12}$ genome copies/ml), AAV2/5-mTH-NLS-CreERT2 (AAV-TH-CreERT2, $5.88\times10^{12}$ genome copies/ml), AAV2/5-mTH-mCherry (AAV-sham, $2.82\times10^{12}$ genome copies/ml), and AAV2/9-hSyn-DIO-ChrimsonR-mCherry ($4.9\times10^{12}$ genome copies/ml) from BrainVTA, Wuhan, China; AAV9-hSyn-DA2m ($4.6\times10^{13}$ genome copies/ml) and AAV9-hSyn-rDA1m ($9.7\times10^{13}$ genome copies/ml) were obtained from Vigene Biosciences, Jinan, Shandong, China; rAAV9-hSyn-rDA3m ($5.88\times10^{12}$ genome copies/ml) were obtained from BrainCase, Shenzhen, China.

In AAV-infected neuron experiments, the following viral vectors were used: AAV2/5-CMV-DIO-Synaptophysin-AT1.03$^{YEMK}$ (Syn-AT1.03$^{YEMK}$, $1.3\times10^{13}$ genome copies/ml) from Obio Technology, Shanghai, China; AAV2/9-EF1a-DIO-4mt-jGCaMP8s (4mt-jGCaMP8s, $3.3\times10^{12}$ genome copies/ml), AAV2/9-CMV-DIO-VMAT2-pHluorin (VMAT2-pHluorin, $2\times10^{13}$ genome copies/ml), and AAV2/9-EF1a-DIO-axon-ATeam1.03$^{YEMK}$ (Axon-AT1.03$^{YEMK}$, $1\times10^{13}$ genome copies/ml) from BrainVTA, Wuhan, China; AAV2/9-hSyn-ChrimsonR-tdTomato (ChrimsonR-tdTomato, $2.2\times10^{13}$ genome copies/ml), AAV2/9-mTH-4mt-RCaMPh (4mt-RCaMPh,

$2\times10^{13}$ genome copies/ml), and AAV2/9-mTH-4mt-Ateam1.03NL (4mt-AT1.03, $2\times10^{13}$ genome copies/ml) from Shanghai Taitool Bioscience, China.

**Stereotaxic surgery**

Mice were anesthetized with 3% inhaled isoflurane in oxygen, placed in a stereotaxic apparatus (RWD Life Science), and maintained at 1.5% isoflurane during surgery. Injection site coordinates were determined using an online brain atlas tool (https://labs.gaidi.ca/mouse-brain-atlas). All AAV viruses were diluted in 1× PBS to a titer range of $3–8\times10^{12}$ genome copies per ml before use. Mitochondrial calcium indicators and optogenetic proteins were selectively expressed in the VTA via unilateral injections of AAV2/9 serotypes carrying DIO-4mt-jGCaMP8s, DIO-GCaMP6m, or DIO-ChrimsonR-mCherry (300 nl, 50 nl/min), targeted to coordinates AP -3.2 mm, ML +0.5 mm, DV -4.2 mm relative to bregma. Dopamine sensors AAV9-hSyn-DA2m, AAV9-hSyn-rDA1m, or rAAV-hSyn-rDA3m were similarly injected into the NAc (100 nl, 50 nl/min, AP +1.6 mm, ML +0.8 mm, DV -4.2 mm). For MCU knockout, AAV-mTH-NLS-Cre, AAV-mTH-NLS-CreERT2 or AAV-mTH-mCherry was bilaterally injected into the VTA following the same parameters. For fiber photometry and optogenetic experiments, optic fibers with a 200-μm diameter core (Thinker Tech) were placed 0.1 mm above the viral injection site. For *in vivo* two-photon imaging, a 1 mm diameter, 4.5 mm long gradient-index (GRIN) lens (GRINTECH NEM-100-25-10-860-S) was implanted into the NAc at coordinates AP +1.6 mm, ML +0.8 mm, DV -4.1 mm, two weeks following AAV injection. Mice were used for experiments no earlier than three weeks post-surgery to ensure adequate expression of AAVs.

**Fiber photometry**

Fiber photometry recording was performed as previously described[1]. Briefly, we implanted an optical fiber above specific brain regions following the injection of various AAVs expressing targeted probes. The fiber was secured to the mouse skull using dental cement. After a three-week expression period, probe fluorescence signals were recorded during behavioral tests. For individual detection of mitochondrial $Ca^{2+}$ or dopamine levels, a fiber photometry system (QAXK-SS-SC-FPS, Thinker Tech) was used. Purple (405 nm) LED light was bandpass filtered (405/10 nm, model 65133, Edmund Optics), blue (473 nm) LED light (Cree LED) was bandpass filtered (470/25 nm, model 65144, Edmund Optics) and yellow (580 nm) LED light (Cree LED) was bandpass filtered (572/28 nm, model 84100, Edmund Optics). These lights were reflected by a dichroic mirror (model 67079，495 nm long pass, Edmund Optics) and another dichroic mirror (model 87-282, multi-band filter, Edmund Optics), and then focused using a 20x objective lens (Olympus). Three types of excitation light were modulated at different frequencies to excite the fluorescent light at distinct wavelengths. An optical fiber guided the light between the commutator and the implanted optical fiber cannula. The excitation light power at the tip of the optical fiber was adjusted to 20-30 μW and delivered at 100 Hz. The green fluorescence was bandpass-filtered (525/39 nm, model MF525-39, Thorlabs), and the red fluorescence was bandpass filtered (615/20 nm, model 87753, Edmund Optics), then collected using two photomultiplier tubes (model H10721-210, Hamamatsu). Signals of different frequencies were demodulated, amplified (C7319, Hamamatsu), and low-pass filtered.

Calcium-dependent fluorescence was elicited by stimulating 4mt-jGCaMP8s or GCaMP6m-expressing neurons and terminals (470 nm, 40 μW; 410 nm, 20 μW for motion artifact correction). Dopamine fluorescence was obtained by stimulating DA2m rDA1m, or rDA3m in the NAc (470 nm or 580 nm, 40 μW).

For concurrent mitochondrial $Ca^{2+}$ imaging and optogenetic stimulation of dopaminergic terminals in the NAc, we co-transfected dopamine neurons with DIO-4mt-jGCaMP8s and DIO-ChrimsonR-mCherry in the VTA of DAT-Cre mice. For dopamine imaging during optogenetic stimulation, we expressed DA2m in the NAc and DIO-ChrimsonR-mCherry in VTA of DAT-Cre mice. We utilized a specialized fiber photometry system (QAXK-SS-SC-FPS-OG, Thinker Tech) for simultaneous optical recording and optogenetic activation. The optical path was designed to combine the optogenetic light and the excitation lights of fiber photometry into a single beam, with the fluorescence of different wavelengths being received separately through dichroic mirrors at the detecting end, minimizing the effect of the optogenetic light on the recording of fluorescence signal recording.

To assess drug-induced changes in mitochondrial calcium transients and dopamine levels, mice underwent a three-day habituation protocol. Subjects were handled and placed in an open field apparatus for 30 minutes daily to acclimate to human interaction and the experimental environment. Subsequently, we recorded mito-$Ca^{2+}$ and dopamine dynamics for a 5-minute baseline period, followed by an intraperitoneal injection of either saline (control), cocaine (20 mg/kg), heroin (10 mg/kg), METH (2 mg/kg), or morphine (30 mg/kg). Recordings continued for 30 minutes post-injection. This protocol allowed for a comprehensive evaluation of acute drug effects on mitochondrial calcium homeostasis and dopamine release across multiple substances of abuse.

For the analysis of mitochondrial calcium and dopamine dynamics during food consumption, we employed a similar initial habituation protocol. Mice were handled and exposed to the open field apparatus for 30 minutes daily over three consecutive days. Prior to the experiment, subjects underwent a 12-hour food and water deprivation period. Mice were then placed the mice in a custom chamber (20×20×22 cm) featuring a central, small platform (1.2 mm diameter, 1 cm height). Dustless precision food pellets (14 mg, Bio-serv) were manually delivered to this platform, one at a time. An overhead infrared camera, synchronized with the optical fiber recording system, captured video footage throughout the session. The onset of each food intake bout was precisely determined through frame-by-frame video analysis, allowing for temporal synchronization between feeding behavior and fiber fluorescence signals.

### Optogenetic stimulation

ChrimsonR activation was achieved using a 589-nm laser system (RWD Life Science, IOS-589) delivering 20 mW at the optical fiber tip (in vivo) or 4 mW from the fiber collimator (in vitro). Light power measurements before and after each session verified system stability. Fixed-frequency stimulation protocols consisted of 7, 15, 30, 60, or

120 pulses at 20 Hz (5 ms square waves, 20 mW), while fixed-pulse protocols delivered 120 pulses at 4, 20, or 40 Hz. Each protocol was repeated five times with 140-second intervals. For each mouse, five repeated stimulation responses were temporally aligned and averaged to generate a mean response curve, which represented the mitochondrial $Ca^{2+}$ or dopamine dynamics for that specific stimulation parameter. These individual mouse response curves were then used for subsequent group analyses. For *in vitro* optogenetic depolarization of dopamine (DA) neurons, a 589-nm laser was used with a 5-second pulse duration at 80 Hz frequency, with 3 ms square wave pulses delivering a power output of 4 mW of light power to the cells from the fiber collimator.

**Photometry data analysis**

The photometry data were analyzed using the Triple Color Analysis Package software (Thinker Tech) based on MATLAB (Version R2021a)[2]. The calcium responses of 4mt-jGCaMP8s and GCaMP6m, elicited by 470 nm excitation, were standardized against the 405/415 nm isosbestic signal. This process employed a linear regression model correlating both signals during the baseline phase, yielding a normalized fluorescence metric, denoted as $F_{normalized}$. This metric represents the fluorescence intensity predicted from the signal captured at 405/415 nm excitation. Data analysis was conducted using the formula $z = \frac{Fnormalized - \mu}{\sigma}$, where $F_{normalized}$ is the normalized photometry signal, $\mu$ signifies the mean $F_{normalized}$ during the pre-stimulus baseline period, and σ is the standard deviation of $F_{normalized}$ within the same baseline interval. The z-score outcomes were used to determine the average z-score values. The values of dopamine signal change ($\Delta F/F_0$) were calculated by $(F_{stimulaton}–F_{baseline})/F_{baseline}$, and $F_0$ is the mean value of the pre-stimulus signal (baseline). For individual traces from mice, both the z-score and the $\Delta F/F_0$ traces underwent smoothing via a moving average filter to enhance data clarity.

Based on the normalized $\Delta F/F_0$ or z-score results, the average z-score was calculated as the mean of z-score values at each time point within the analysis period. The Area Under Curve (AUC) was calculated by summing the $\Delta F/F_0$ or z-score results during the indicated time after stimulation. Peak values were determined as the maximum $\Delta F/F_0$ or z-score within the stimulation or analysis period. For reward-related signals, analysis windows spanned 35 minutes or 6 minutes, with 5-minute or 1-minute pre-stimulus baselines, respectively. For optically evoked responses, a 32-second analysis window was used with a 2-second pre-stimulus baseline.

**In vivo two-photon imaging**

To visualize mitochondrial $Ca^{2+}$ dynamics in dopaminergic axon terminals within the nucleus accumbens (NAc), we employed an advanced fast, high-resolution miniaturized two-photon microscope (FHIRM-TPM V2.0) for *in vivo* imaging. The experiments were conducted at the Nanjing Brain Observatory (PKU-Nanjing Joint Institute of Translational Medicine). For this study, we injected 300 nL of AAV2/9-DIO-4mt-jGCaMP8s into the ventral tegmental area (VTA; AP: -3.2, ML: 0.5, DV: -4.2) of the DAT-Cre mice. Following two weeks of viral expression, we created a 4 mm diameter cranial window above the injection site using a miniature hand-held drill (RWD Life Science). A 4 mm glass coverslip (100 μm thickness) was inserted into the window and secured with biological tissue glue (3M, USA). After a recovery period of at least one week, an imaging baseplate was affixed over the cranial window and cemented with dental acrylic. The switchable FHIRM-TPM V2.0 platform, featuring a high-resolution mode (field of view: 190×190 $\mu m^2$; resolution: ~850 nm; working distance: 400 μm) and a large-field mode (field of view: 420×420 $\mu m^2$; resolution: ~1300 nm; working distance: 1000 μm), was mounted onto a permanently fixed baseplate over the cranial window. Initially, fluorescent regions of interest (ROIs) were identified using a benchtop microscope, after which the high-resolution miniaturized two-photon microscope (mini-2P) was mounted for targeted imaging.

To secure the setup, a headpiece was affixed to the baseplate following craniotomy, and 1.5% low-melting-point agarose was applied to cover the chronic window prior to imaging. Mice were habituated for one-week post-headpiece implantation, with the headpiece being protected between sessions using parafilm and plasticine to maintain window cleanliness. Before each imaging session, the cover of the holder and the protective glue (Kwik-Cast, WPI Inc., USA) over the cranial window were carefully removed. The headpiece was then mounted onto the holder and secured with M2 screws. Imaging was performed at a frame rate of 9.62 Hz (512×512 pixels) using a femtosecond fiber laser (~35 mW at the objective, TVS-FL-01, Transcend Vivoscope Biotech Co., Ltd, China), with 4mt-jGCaMP8s being excited at 920 nm. Data acquisition was managed using GINKGO-MTPM software (Transcend Vivoscope Biotech Co., Ltd, China).

**Primary midbrain neuron culture**

*DAT-Cre; MCU*$^{+/+}$ (WT) and *DAT-Cre: MCU*$^{flox/flox}$ (DAT-*MCU* cKO) midbrain neurons were cultured from E12.5 embryonic mice of random sex. Genotyping was performed to separate WT and cKO midbrain tissue. Neurons were dissociated using 0.25% Trypsin-EDTA solution (Merck, T4049) and plated onto poly-D-lysine (Thermo Fisher; A3890401) coated 8-chamber Lab-Tek II chambered #1.5 German coverglass systems (Nunc,155411) in plating medium (high glucose DMEM supplemented with 10% FBS, 10% Horse Serum, and 1% Penicillin-Streptomycin) (all purchase from Thermo Fisher Scientific) ($2.5\times10^5$ cell/per well).

After 6 hours, the plating medium was replaced with neuronal precursor induction medium (Neurobasal A medium supplemented with 1% B-27 Plus, 0.5 mM GlutaMAX, 1% N-2 Supplement, 1% Penicillin-Streptomycin, 10 ng/ml EGF, 10 ng/ml FGF) (all purchased from Thermo Fisher Scientific) until cultured for two days *in vitro* (DIV2). Subsequently, neurons were cultured in neuronal differentiation medium (Neurobasal A medium supplemented with 1% B-27 Plus, Dopaminergic Neuron Maturation Supplement (50×, Gibco, A3147401), 1% Penicillin-Streptomycin), with half of the medium replaced every two days. Neurons were transfected with various constructs at DIV4-6 using AAV infection and imaged at DIV9-12 using a GE DeltaVision microscope.

### Immunostaining for dopamine neuron verification

Neurons used in the experiments were verified as dopamine neurons through post-experiment immunostaining for tyrosine hydroxylase (TH). Cultures were fixed for 15 minutes with 4% paraformaldehyde in PBS, followed by permeabilization with 0.2% Triton X-100 for 10 minutes. Blocking was performed with 3% normal goat serum (NGS) in PBS for 1 hour. Cells were then incubated at room temperature for 1 hour with primary antibodies: mouse anti-MAP2 monoclonal antibody (Abcam, ab11268) at 1:200 and rabbit anti-TH polyclonal antibody (Abcam, ab112) at 1:200 for 1 hour. Cross-adsorbed secondary antibodies from Molecular Probes were subsequently applied: Alexa Fluor 488-conjugated goat anti-rabbit IgG and Alexa Fluor 594-conjugated goat anti-mouse IgG, both at a 1:1000 dilution, and incubated for 1 hour at room temperature in the dark.

### Viral-mediated gene delivery and fluorescent probes expression

We employed various viral-mediated gene delivery approaches to express specific probes and optogenetic proteins in neurons cultured *in vitro*. The isolated neurons were planted at $2.5\times10^5$ cells per well in an 8-chamber coverglass system. For presynaptic vesicle release detection, AAV2/9-DIO-VMAT2-pHluorin ($2\times10^{10}$ genome copies/well) was used to express VMAT2-pHluorin, enabling the monitoring of monoamine neurotransmitter dynamics within synaptic vesicles. To detect KCl-induced mitochondrial calcium influx and changes in mitochondrial ATP levels, AAV2/9-mTH-4mt-RCaMPh ($2\times10^{10}$ genome copies/well) and AAV2/9-mTH-4mt-Ateam1.03NL ($2\times10^{10}$ genome copies/well) were introduced into both WT and DAT-MCU cKO neurons, expressing the red mitochondrial calcium indicator 4mt-RCAMPh and the mitochondrial ATP sensor 4mt-AT1.03 under the tyrosine hydroxylase (TH) promoter. Presynaptic ATP detection was achieved by co-expressing AAV2/5-CMV-DIO-Synaptophysin-AT1.03$^{YEMK}$ ($4\times10^{10}$ genome copies/well) and AAV2/9-hSyn-ChrimsonR-tdTomato ($2.2\times10^{10}$ genome copies/well) in both WT and DAT-*MCU* cKO dopamine neurons, while AAV2/9-EF1a-DIO-axon-ATeam1.03$^{YEMK}$ ($1\times10^{10}$ genome copies/well) was utilized for axonal ATP detection. For cytoplasmic calcium changes, Fluo4 AM (2 μM) or Calbryte™ 590 AM (20 μM) were used to stain mature dopaminergic neurons at DIV9 for 30 minutes before imaging optical or KCl-induced changes. To detect KCl-induced mitochondrial calcium influx under Berberine treatment, AAV2/9-EF1a-DIO-jGCaMP8s ($1.5\times10^{10}$ genome copies/well) was introduced into dopamine neurons.

**Imaging mitochondrial calcium influx and ATP dynamics**

Infected neurons were transferred to pre-warmed Tyrode's solution (119 mM NaCl, 2.5 mM KCl, 50 mM HEPES (pH 7.4), 2 mM $CaCl_2$, 2 mM $MgCl_2$, 10 mM glucose), supplemented with 10 μM 6-cyano-7-nitroquinoxaline-2,3-dione (CNQX) and 50 μM D, L-2-amino-5-phosphonovaleric acid (APV). The temperature was maintained at 37°C using an air stream incubator. Imaging was performed using a GE DeltaVision microscope equipped with a 40×1.3 NA oil immersion or 20×0.8 NA objective. Time-lapse images were acquired at a 1024×1024-pixel resolution.

For mitochondrial calcium influx and ATP imaging, images were captured at 1-minute intervals over a 20-minute period. Mitochondrial, presynaptic, and axonal ATP levels were measured using neurons expressing 4mt-AT1.03, Syn-AT1.03$^{YEMK}$, or Axon-AT1.03$^{YEMK}$, respectively. ATeam was excited at 435 ± 5 nm and the ratio of emission at 527 nm (YFP) and 475 nm (CFP) was plotted. Ratiometric images were generated using ImageJ (NIH).

To assess presynaptic ATP levels or cytoplasmic calcium changes under repeated stimuli, neurons were kept in modified Tyrode's solution (119 mM NaCl, 2.5 mM KCl, 50 mM HEPES (pH 7.4), 2 mM $CaCl_2$, 2 mM $MgCl_2$, 1.25 mM lactate and 1.25 mM pyruvate) supplemented with 10 μM CNQX and 50 μM APV, and stimulated with 400 pulses (80 Hz, 3 ms pulse width, 5 s, 4 mW light from the fiber collimator ) delivered via a 589 nm optogenetics system (RWD Life Science, IOS-589) coupled with a fiber collimator (QAXK-JCQ, Thinker Tech Nanjing Bioscience Inc.). Images were initially recorded for 3 minutes as a baseline, and further captures were made 7 minutes after a 5-second optical stimulation, with 1-minute intervals.

For axonal ATP measurements, dopaminergic neurons at DIV10 expressing Axon-AT1.03$^{YEMK}$ were imaged using a 20× 0.8 NA objective. After 30 minutes of baseline imaging, ATP (5 mM) or phosphate-buffered saline (PBS) was added to the culture medium, and subsequent images were taken at 10-minute intervals for 150 minutes. KCl-induced mitochondrial calcium influx and ATP changes were detected in DIV9 dopaminergic neurons expressing 4mt-RCaMPh and 4mt-AT1.03. Images were captured at 1-minute intervals for 20 minutes. To investigate the effect of berberine on KCl-induced mitochondrial calcium influx, DIV9 dopamine neurons (from *DAT-Cre: MCU*$^{+/+}$ *mice*) expressing AAV-DIO-4mt-jGCaMP8s were imaged every 10 seconds for 250 seconds, with 75 mM KCl added after 50 seconds of baseline imaging.

**Presynaptic vesicle release analysis**

To examine presynaptic vesicle release under KCl treatment, differentiated dopamine neurons were infected with AAV-DIO-VMAT2-pHluorin at DIV7. Experiments were conducted at DIV9-10 in Tyrode's solution supplemented with 10 µM CNQX and 50 µM APV, maintained at 37°C. Using a GE DeltaVision microscope with a 40× 1.3 NA objective, baseline fluorescence was recorded for one minute, followed by image capture every 3 seconds after 75 mM KCl stimulation. The exposure time for fluorescein isothiocyanate (FITC) was set to 50 ms.

The total pool (TP) of synaptic vesicles was identified by perfusion with Tyrode's alkaline solution containing 50 mM $NH_4Cl$ (pH 7.4). Fluorescence data were analyzed using ImageJ with Time Series Analyzer plug-ins. Puncta exhibiting no lateral movement during image acquisition and responding to KCl stimulation were selected for analysis. Background correction was performed by subtracting the intensity of a nearby cell-free region from the imaged cell signal.

Changes in fluorescence intensity were calculated using the formula $\Delta F/F_0 = (F-F_0)/F_0$, where $F_0$ represents the initial, background-subtracted fluorescence intensity, and F represents the background-subtracted fluorescence intensity at subsequent time points. Data were then normalized to the peak fluorescence observed during $NH_4Cl$ stimulation and expressed as a percentage of the total pool. Each experimental group involved observations of at least six dopamine neuron cells, with a minimum of 15 boutons selected per cell. The resulting plots represent the mean ± S.E.M. of normalized $\Delta F/F_0$ curves for each cell. For quantitative analysis of differences between groups, the peak VMAT2-pHluorin fluorescence for each cell was normalized to the total pool following KCl stimulation. The mean ± S.E.M. of these normalized values was then used for subsequent statistical analyses.

**Immunoblot**

Bilateral midbrain and striatum samples were extracted from the mouse brains and homogenized in lysis buffer (320 mM sucrose, 4 mM HEPES pH 7.4, 1 mM $MgCl_2$, 0.5 mM $CaCl_2$, 5 mM NaF, EDTA-free protease inhibitor cocktail) on ice. Proteins were separated by SDS-PAGE and transferred to PVDF membranes. Membranes were incubated with primary antibodies overnight at 4°C, followed by HRP-conjugated secondary antibodies (Santa Cruz, 1:5000). The primary antibodies were used: Rabbit anti-MCU (1:1000, CST, 14997), Rabbit anti-TH (1:2000, Abcam, ab112), Mouse anti-DAT (1:2000, Abcam, ab128848), and Mouse anti-α-tubulin (1:5000, Sigma-Aldrich, T5168).

**Immunohistochemistry and microscopic analysis**

Paraffin multiplex immunohistochemistry (mIHC) was employed to validate the conditional knockout of MCU. For other immunohistochemical analyses, frozen section IHC was used. Mice were transcardially perfused with 20 mL 0.01 M phosphate buffered saline (PBS, pH 7.4) followed by 20 mL of 4% paraformaldehyde solution (PFA) diluted in PBS (pH 7.4) under deep isoflurane anesthesia. To validate the expression of 4mt-jGCAMP8s and GCAMP6m, the perfusion solution was comprised of 15 mL of PBS supplemented with 10 mM $CaCl_2$. The brains were then dehydrated, embedded in paraffin, and sectioned into 3-μm paraffin slices. Alternatively, for frozen sections, after gradient sucrose dehydration, brains were embedded in OCT compound and sectioned into 8-μm frozen sections.

To confirm the conditional knockout of MCU, multiplex immunohistochemistry was employed to label tyrosine hydroxylase (TH) and MCU. Sections were deparaffinized using a graded ethanol series and underwent antigen retrieval in Tris-EDTA buffer (pH=9) with heat. After cooling, the sections were incubated with 3% hydrogen peroxide at room temperature for 15 minutes to quench endogenous peroxidase activity. Non-specific binding was blocked with a blocking solution for 30 minutes at room temperature. Sections were incubated with primary antibodies overnight at 4 ° C, followed by incubation with corresponding secondary antibodies at room temperature for 30 minutes. After washing with PBS, the sections were stained with DendronFluor TSA (Histova, NEON 4-Color IHC Kit for FFPE, diluted 1:100, for 30-60 seconds). The AbCracker® wax section repair/rapid removal dual-purpose buffer was used to thoroughly remove the primary and secondary antibodies. Antigens were labeled with different fluorophores in a sequential manner. Finally, DAPI was used to stain the nuclei. The primary antibodies were: Rabbit anti-MCU(1:2000, CST, 14997), Rabbit anti-TH(1:2000, Abcam, ab112).

For other antigens labeling, immunofluorescence was performed as previously described[3]. Sections were washed in PBS and incubated with 1% bovine serum albumin (BSA) and 0.3% Triton X-100 for 15 minutes. After incubation with 5% normal goat serum (NGS)/PBS was added for 30 minutes, primary antibodies were applied overnight at 4°C in PBS containing 1% BSA and 0.3% Triton X-100. Sections were then washed and incubated with secondary antibodies for 2 hours at room temperature. Following antibody detection, sections were finally stained with DAPI. The primary antibodies used were as follows: Chicken anti-GFP(1:400, Invitrogen, A10262), Rabbit anti-TH(1:400, Abcam, ab112), Mouse anti-COXIV(1:400, CST, 11967S), Rabbit anti-mCherry(1:400, Abcam, ab167453), Rabbit anti-p-TH(ser40) (1:400, GeneTEX, GTX16557). The stained brain sections were subsequently examined using a high-resolution Olympus SLIDEVIEW$^{TM}$ VS200 microscope (Olympus, Germany), and quantitative image analysis was performed using ImageJ software suite.

**Drug-induced locomotor activity assays**

Locomotor activity was assessed using the LabState Vision system (AniLab Scientific Instruments). Drug-naïve mice were acclimated to an open-field arena (35 cm × 35 cm) for 60 minutes before receiving intraperitoneal (i.p.) injections of 10 mg/kg heroin, 30 mg/kg morphine, or 2 mg/kg methamphetamine (METH). Locomotor activity was recorded for 3 hours post-injection to evaluate the behavioral response. In the cocaine-induced locomotor activity test, mice underwent a 30-minute adaptation period to the test environment before receiving an i.p injection of 20 mg/kg cocaine. Their locomotor activity was then recorded for 1 hour to capture the acute effects of cocaine. To investigate the impact of berberine on heroin-induced locomotor activity, mice were first adapted to the open field for 1 hour. They then received an intra- NAc infusion of berberine. After a 30-minute rest period, they were administered a 10 mg/kg heroin injection. Locomotor activity was recorded for 3 hours to evaluate berberine's modulatory effects on heroin-induced behavior.行为敏化没写

**Conditioned place preference (CPP)**

The CPP procedure was conducted in a standard two-chamber CPP apparatus (30 cm × 25 cm × 20 cm) inside a soundproof box (AniLab, Ningbo, China). Time spent in each chamber was recorded and analyzed using infrared beams and an automated analysis system during both baseline and test periods, as previously described[4]. The CPP test followed a standard method with minor modifications. The procedure included three main phases: pre-test, conditioning, and post-test.

During the pre-test phase, the mice had free access to both chambers with the gate open, and their baseline movement over 1800 seconds was recorded to identify initial preferences. Mice with strong initial preferences (spending disproportionately more or less time on one side) were excluded. The remaining mice were randomly assigned to experimental groups. For the conditioning phase, the guillotine door was closed, confining the animals to a specific compartment for 30 minutes each day for eight days. Mice were given intraperitoneal injections of heroin (3 mg/kg), cocaine (10 mg/kg), or METH (2 mg/kg) on alternating days, with the drug-paired compartment used on the corresponding day. On alternate days, they received saline injections and were placed in the saline-paired compartment. In the post-test phase, the gate separating the two chambers was removed to allow unrestricted movement. Mice were randomly placed in one chamber, and the time spent in each chamber was recorded for 1800 seconds. The CPP score was determined by calculating the difference in time spent in the drug-paired versus the saline-paired compartment, expressed in seconds.

For food-based CPP, the procedure followed the same protocol as the drug-induced CPP, except during the conditioning phase. Mice underwent a 5-day training period, with morning sessions involving milk tablets exposure in one box for 50 minutes, and afternoon sessions in a different box without milk tablets for another 50 minutes. Training was conducted twice daily, with a 6-hour interval between sessions.

**Heroin CPP-induced reinstatement test**

The heroin CPP-induced reinstatement test was performed as previously described methods, with slight modifications[5]. The modified experimental procedure comprised five phases: pre-test, conditioning, post-test, extinction, and reinstatement. The first three phases followed the protocol described above. During the extinction phase, the mice were randomly placed in one chamber and allowed to explore both chambers freely for 30 minutes. The time spent in each chamber was recorded daily and the CPP score was calculated. Once the CPP score approached zero, indicating extinction of the conditioned preference, the next phase was initiated. For the heroin-induced reinstatement test, mice first received an intra-NAc infusion of berberine. Thirty minutes later, they were intraperitoneally injected with 1.5 mg/kg of heroin and then subjected to a preference test. During this test, the time spent in each chamber was recorded over an 1800-second period of free movement.

**Cocaine self-administration paradigm**

Surgical procedures were performed under isoflurane anesthesia (3% induction, 1-1.5% maintenance). A Micro-Renathane catheter (MRE-025, 5.5 cm, Braintree Scientific) was advanced 1.2 cm into the right jugular vein and secured to a right-angled 24-gauge steel cannula (AniLab, Ningbo, China). The assembly was anchored to the skull using cyanoacrylate adhesive and dental cement. To maintain catheter patency and prevent infection, catheters were flushed daily with 0.05 ml saline containing heparin (20 IU/ml) and penicillin sodium (8,000 units/ml).

Following post-surgical recovery (5-7 days), mice underwent behavioral training in operant chambers (AES-DSA, AniLab) equipped with dual nose-poke ports positioned 2 cm above floor level. Each chamber featured a cue light mounted 5 cm above the active port and ambient illumination maintained throughout the 2-h daily sessions. Responses at the active port triggered intravenous cocaine delivery (0.5 mg/kg/infusion; 0.015 mL over 0.75 s) coupled with a 5-s light cue presentation, followed by a 4.25-s timeout period during which additional responses were recorded but had no programmed consequences. Responses at the inactive port were monitored but produced no outcomes.

To facilitate acquisition of drug-seeking behavior, 5-6 experimenter-delivered cocaine infusions were administered over 5-10 minutes during the first two training days (data excluded from analyses). Daily sessions were capped at 50 infusions until stable self-administration patterns emerged, defined by three criteria: (i) consistent procurement of >20 infusions per session across three consecutive days, (ii) less than 20% variability in daily infusion numbers between consecutive sessions, and (iii) discrimination between active and inactive ports reflected by a response ratio exceeding 2:1.

Upon meeting acquisition criteria, the mice underwent extinction training followed by cue-induced reinstatement testing in the same operant chambers. During the extinction phase, responses at either port were recorded but produced no programmed consequences. In reinstatement tests, active port responses triggered presentation of the previously drug-paired light cue without cocaine delivery. For pharmacological intervention studies examining berberine's effects on cue-induced reinstatement, 3 mg/kg berberine or vehicle (ultra-pure water) was administered systemically 2 h prior to reinstatement testing.

**Dopamine measurement of mouse nucleus accumbens**

Dopamine concentrations in the nucleus accumbens were measured using high-performance liquid chromatography (HPLC) coupled with electrochemical detection. Bilateral samples of the nucleus accumbens were extracted from the mouse brain and homogenized in centrifuge tubes containing grinding beads. Each sample was treated with 200 μL of freshly prepared 0.1 M HCl solution, and homogenization was conducted at 5200 rpm for 20 seconds. The homogenate was centrifuged at 12,000×g for 10 minutes at 4°C, and the supernatant was harvested and passed through a 0.22μm PVDF syringe filter.

The HPLC mobile phase, prepared with 100 mM sodium dihydrogen phosphate monohydrate, 0.74 mM 1-octanesulfonic acid sodium salt, 34 μM EDTA, and 10% methanol, adjusted to pH 3.0, was filtered and degassed before use. It was delivered at a flow rate of 0.6 mL/min via an HPLC pump (S 1130 HPLC pump system, Sykam). Dopamine and its metabolites were resolved chromatographically and quantified using a C18 HPLC column (2.1×100 mm, 3 μm, Acutec Scientific) and an electrochemical detector (DECADE Lite, Acutec Scientific). Dopamine was identified based on retention times compared to authentic standards. Sample concentrations were calculated using standard curves generated from known concentrations of dopamine. Data were normalized relative to the total wet weight of the tissue samples.

**Intra-NAc drug infusion**

Thirty minutes before behavioral experiments or fiber photometry, mice received an intra-NAc infusion of berberine (0.3 μg per side) or an equivalent volume of vehicle. The infusion was delivered via an injector cannula using a micro-infusion pump (RWD Life Science) at a steady rate of 0.1 μL/min, culminating in a total volume of 1 μL per hemisphere. After infusion5min, the injector cannula were carefully removed, and the mice were allowed to recuperate for 1 hour before proceeding with behavioral assessments. This protocol ensured adequate drug distribution and pharmacological stabilization for subsequent analyses.

**Rotarod test**

Motor coordination and balance were assessed using a rotarod apparatus. On the day before testing, mice underwent a habituation trial, where they were placed on a five-lane rotarod treadmill (Ugo Basile) operating at a constant speed of 4 rpm for up to 300 seconds. From days 1 to 4 of testing, mice underwent three consecutive trials per day with 1-hour inter-trial intervals. The rotarod speed accelerated from 4 to 40 rpm over 300 seconds, and the latency to fall off the rod was recorded.

**Novel object recognition test**

Recognition memory was assessed using the novel object recognition test. During the habituation phase, mice were placed in a Plexiglas arena (50 × 50 × 50 cm) for 5 minutes. Three hours later, during the acquisition phase, two identical objects were placed in opposite corners of the arena, and mice explored them for 10 minutes. After 24 hours, one of the objects was replaced with a novel object, and mice explored the arena for another 10 minutes. The novel object recognition index was calculated as the ratio of time spent with the novel object to the total time spent with both objects. Object interactions were defined as the mouse's nose being within 1 cm of the object, including touching and sniffing. Rearing on objects was not scored. Behavior was recorded and analyzed using the VisuTrack Animal Behavior Analysis System (XinRuan Co. Ltd, Shanghai).

**Y-maze spontaneous alternation test**

Spatial cognition was evaluated using Y-maze spontaneous alternation test. The Y-maze consisted of three arms (30 cm length × 5 cm width × 15 cm height) intersecting at 120° angles. Mice were placed at the center and allowed to explore for 5 minutes. An entry was counted when the mouse's hindpaws completely entered an arm. Correct alternations occurred when the mouse entered three different arms consecutively. The spontaneous alternation percentage was calculated as: Spontaneous Alternation (%) = Alternations/ (Arm Entries−2) ×100.

**Open field Test (OFT)**

The open-field test was conducted to evaluate anxiety and motor ability. Mice were placed in a square open-field arena (50 cm × 50 cm) and allowed to move freely for 10 minutes (anxiety assessment) or 30 minutes (motor ability assessment). Behavior was recorded using the VisuTrack Animal Behavior Analysis System (XinRuan Co. Ltd,

Shanghai), which tracked the distance traveled, number of entries, and time spent in the center versus the periphery zones.

**Elevated Plus Maze Test (EPM)**

The elevated plus maze test is commonly used to evaluate anxiety-like behaviors in mice. The apparatus consists of two closed arms (25 cm × 5 cm × 16 cm) opposite each other and perpendicular to two open arms (25 cm × 5 cm × 0.5 cm), with a central platform (5 cm × 5 cm × 0.5 cm). In the open arms, a small wall (0.5 cm high) is used to reduce the risk of falls. The entire apparatus is elevated 50 cm above the floor. At the start of each session, an animal was placed in the central square facing the open arms and allowed to explore the environment freely for 5 minutes. To eliminate olfactory cues from the previous animal, the apparatus was cleaned with 70% ethanol after each test. Performance was recorded by a video camera mounted 100 cm above the maze using the VisuTrack Animal Behavior Analysis System (XinRuan Co. Ltd, Shanghai).

**Forced swim test (FST)**

The forced swim test is a classic behavioral assay used to evaluate despair-like behaviors indicative of depression. Mice were acclimated to the test room for three hours before behavioral testing. Individually, mice were placed in an acrylic cylinder (diameter, 15 cm; height, 25 cm) filled with water (23–25°C) and observed swimming for 6 minutes under normal light conditions. The water depth was adjusted to prevent the animals from touching the bottom with their tails or hind limbs. The immobility time during the last 4 minutes of the 6-minute test was recorded using the VisuTrack Animal Behavior Analysis System (XinRuan Co. Ltd, Shanghai). Following the test, the mice were dried with a paper towel and returned to their home cage.

**Tail suspension test (TST)**

The tail suspension test was utilized to induce a state of despair and helplessness in mice for the evaluation of depression-like behaviors. The test was performed using a specially manufactured tail-suspension box (30 × 30 × 60 cm). Within the box, a medium binder clip hung 50 cm from the base. The tail of each mouse was taped to this clip, and the mice were suspended for 6 minutes. The immobility time during the last 4 minutes of the 6-minute test was recorded using the VisuTrack Animal Behavior Analysis System (XinRuan Co. Ltd, Shanghai).

**Sucrose preference test (SPT)**

The sucrose preference test is a widely recognized method for evaluating anhedonic behavior, a core symptom of depression. Mice were trained for two days to choose between drinking water and a 1% sucrose solution presented in two separate bottles, followed by a 24-hour food and water deprivation and a 24-hour testing phase. On the day of testing, two pre-weighted bottles, one containing 1% (w/v) sucrose solution and the other potable water, were presented to each mouse at the dark phase (20:00). To control side preference in drinking behavior, the position of the bottles was switched after 12 hours of training or testing. Fluid consumption was measured by weighing the bottles before and after the test period. Sucrose preference was calculated as (sucrose intake / (sucrose intake + water intake)) × 100 and expressed as a percentage.

**Morris water maze (MWM)**

The Morris Water Maze (MWM) experiment was executed with modifications to the standard protocol[6]. For the training phase, a transparent circular escape platform (diameter, 10 cm) was submerged 0.5 to 1 cm beneath the surface of water made opaque with non-toxic white paint and consistently positioned in the center of the target quadrant. Initially, mice were permitted a 10-second acclimatization period on the platform. Following this, they were carefully introduced into the maze facing the pool wall and allowed 60 seconds to locate the platform. If the mice did not locate the platform within this time, they were gently guided to it. After each trial, mice remained on the platform for 20 seconds before being re-introduced from a different starting point. This sequence was repeated for a total of four trials. After the trials, the mice were dried with a paper towel and returned to their home cages. 24 hours after the initial training, the mice underwent another training session following the same protocol but without initial acclimatization. This training regimen continued for five consecutive days. On the final day, after the fourth trial, the mice were given a rest period in their home cages. Subsequently, they were subjected to a 60 second probe trial in the water maze without the platform to evaluate spatial memory. Behavioral recordings were captured on video and analyzed using VisuTrack Animal Behavior Analysis System (XinRuan Co. Ltd, Shanghai). The Morris Water Maze was conceptually divided into quadrants, with the target quadrant being the location of the submerged platform. The escape latency, the time taken to find the platform, was monitored throughout the training period as a measure of spatial learning and memory efficacy.

**Statistical analysis**

Statistical analyses and graphical representations were performed using Prism 9.5.0 (GraphPad Software), except for fiber photometry analysis, which is described in detail above. Distribution normality was assessed using D'Agostino-Pearson or Shapiro-Wilk normality test. Experiments containing two groups were analyzed using a two-tailed paired or unpaired Student's t-test for normally distributed data and using a Wilcoxon matched-pairs signed-rank test or Mann-Whitney test for non-normally distributed data. Experiments involving more than two groups were analyzed by one-way ANOVA followed by Bonferroni multiple comparison's test for normally distributed data or Kruskal-Wallis test followed by Dunn's multiple comparison's test for non-normally distributed tests. Data grouped based on more than one nominal variable were analyzed using a two-way ANOVA followed by Bonferroni post hoc test. Correlation was analyzed using the Pearson correlation coefficient test. All statistical details are provided in the Supplementary Table 1. Results are presented as means ± standard errors of the means (SEM). Significance was set as $^{*}P < 0.05$, $^{**}P < 0.01$, and $^{***}P < 0.001$ for all data. The sample size (*n*) refers to the number of cells or animals included in the Figure legends. Mice were randomly assigned to treatments and conditions, and investigators were blinded to genotypes and behavioral outcomes. All the experiments were reliably reproduced for two to three batches.

**References**


1 Zhou, K. *et al.* Reward and aversion processing by input-defined parallel nucleus accumbens circuits in mice. *Nat Commun* **13**, 6244, doi:10.1038/s41467-022-33843-3 (2022).

2 Dong, H. *et al.* Genetically encoded sensors for measuring histamine release both in vitro and in vivo. *Neuron* **111**, 1564-1576 e1566, doi:10.1016/j.neuron.2023.02.024 (2023).

3 Chen, J. *et al.* Glioblastoma stem cell-specific histamine secretion drives pro-angiogenic tumor microenvironment remodeling. *Cell Stem Cell* **29**, 1531-1546 e1537, doi:10.1016/j.stem.2022.09.009 (2022).

4 Sjulson, L., Peyrache, A., Cumpelik, A., Cassataro, D. & Buzsaki, G. Cocaine Place Conditioning Strengthens Location-Specific Hippocampal Coupling to the Nucleus Accumbens. *Neuron* **98**, 926-934 e925, doi:10.1016/j.neuron.2018.04.015 (2018).

5 Yan, P. *et al.* Hypoxia-inducible factor upregulation by roxadustat attenuates drug reward by altering brain iron homoeostasis. *Signal Transduct Target Ther* **8**, 355, doi:10.1038/s41392-023-01578-2 (2023).

6 Zhou, W. *et al.* Loss of function of NCOR1 and NCOR2 impairs memory through a novel GABAergic hypothalamus-CA3 projection. *Nat Neurosci* **22**, 205-217, doi:10.1038/s41593-018-0311-1 (2019).
