## Supplementary Figures for "Mitochondrial calcium influx-driven bioenergetics in the dopaminergic system selectively enable drug reward"

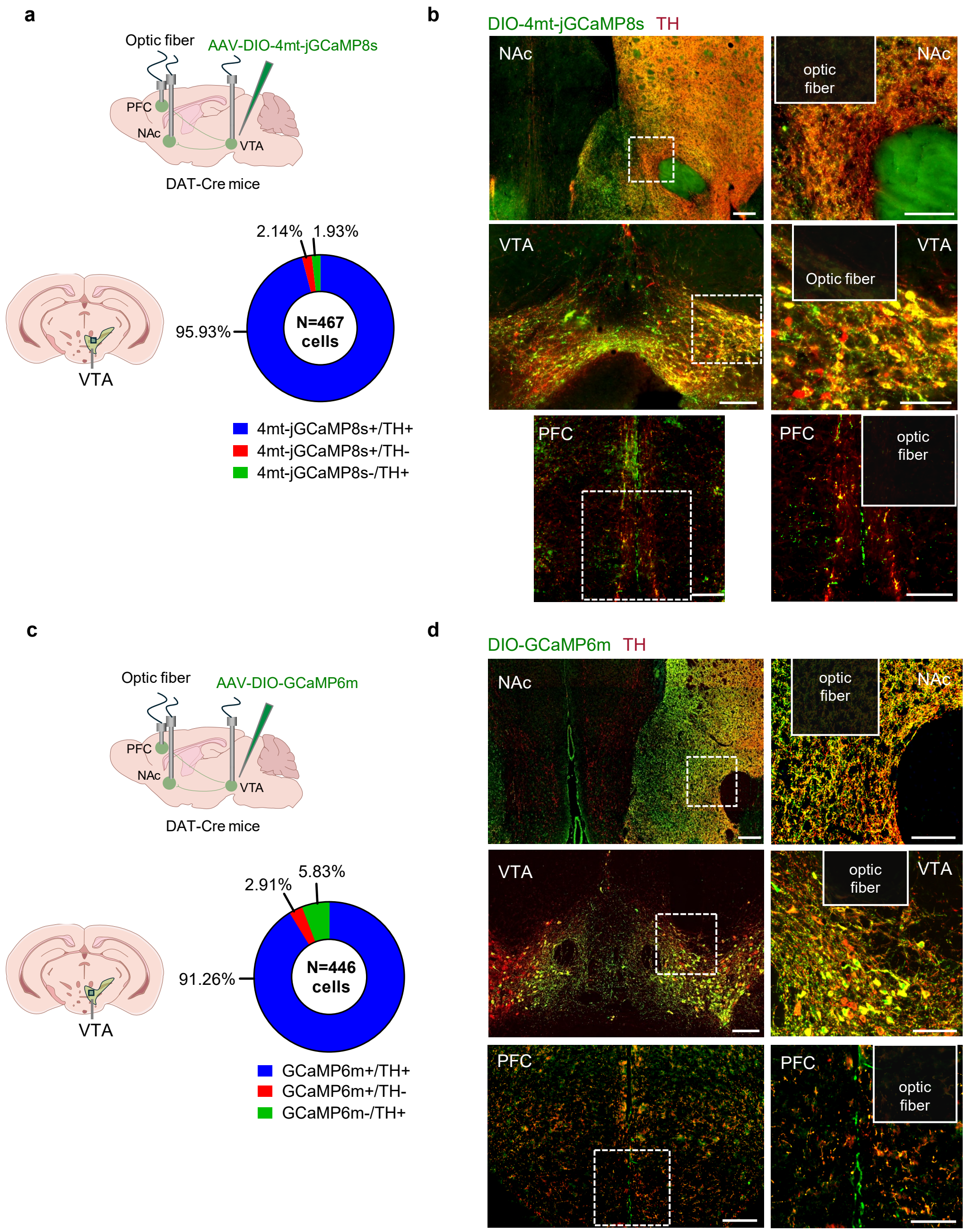


**Extended Data Figure 1. Specific targeting of mitochondrial $Ca^{2+}$ and cytosolic $Ca^{2+}$ probes to $TH^{+}$ neurons in DAT-Cre mice.**

**a**, (Top) Schematic of unilateral stereotaxic injection of AAV-DIO-4mt-jGCaMP8s into the VTA and fiber implantation in VTA, NAc, and PFC. (Bottom) Pie chart showing the co-expression of 4mt-jGCaMP8s with $TH^{+}$ neurons in the VTA and its specificity to $TH^{+}$ neurons.

**b**, (Left) Representative images of 4mt-jGCaMP8s (green) expression in $TH^{+}$ neuron (red) in the VTA, NAc, and PFC. Scale bar: 200 µm. (Right) Magnified view showing optic fiber implants. Scale bar: 100 µm.

**c**, (Top) Schematic of unilateral stereotaxic injection of AAV-DIO-GCaMP6m into the VTA and fiber implants in VTA, NAc, and PFC. (Bottom) Pie chart showing the co-expression of GCaMP6m with $TH^{+}$ neurons in the VTA and its specificity to $TH^{+}$ neurons.

**d**, (Left) Representative images of GCaMP6m (green) expression in $TH^{+}$ neuron (red) in the VTA, NAc, and PFC. Scale bar: 200 µm. (Right) Magnified view showing optic fiber implants. Scale bar: 100 µm.

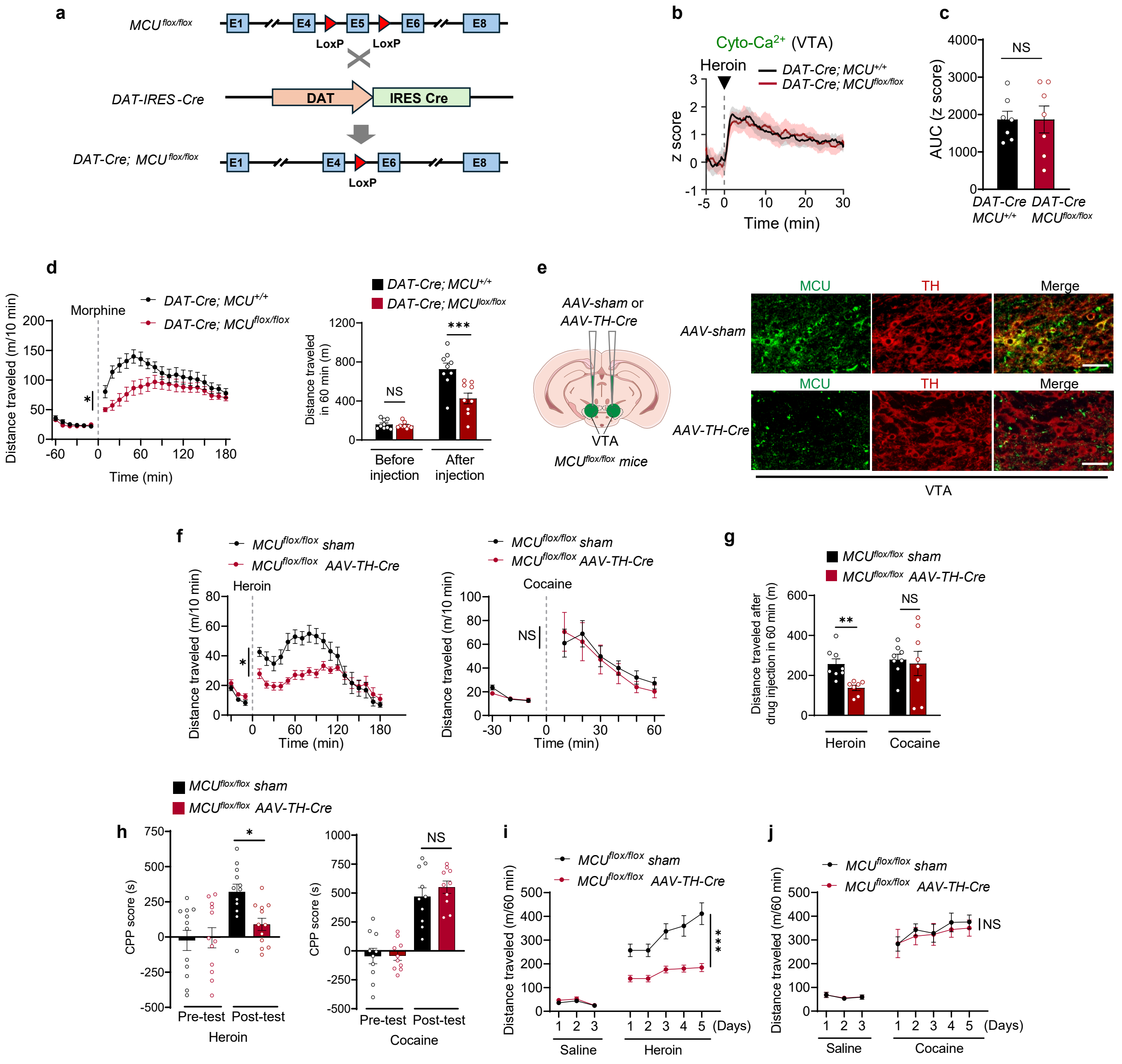

**Extended Data Figure 2. Dopaminergic neuron-specific MCU depletion impairs reward behaviors induced by opioid but not by cocaine.**

**a**, Genetic strategy for selectively delete *MCU* from dopamine transporter (DAT)-positive neurons in the brain.

**b**,**c**, Mean fluorescence traces (**b**) and area under the curve (AUC) values (**c**) showing cytoplasmic calcium responses in VTA dopaminergic neurons following a 10 mg/kg heroin intraperitoneal injection in *DAT-Cre; MCU*$^{+/+}$ and *DAT-Cre; MCU*$^{flox/flox}$ mice ($n$ = 7 per group).

**b**, Traces of cytoplasmic $Ca^{2+}$ dynamics (GCaMP6m fluorescence, z score) in VTA dopaminergic neurons aligned to heroin injection (grey dotted line and arrow) in *DAT-Cre; MCU*$^{+/+}$ (WT) and *DAT-Cre; MCU*$^{flox/flox}$ (DAT-*MCU* cKO) mice ($n$ = 7 per group).

**c**, Bar graph showing area under the curve (AUC) of cytoplasmic $Ca^{2+}$ signals (GCaMP6m fluorescence) in VTA dopaminergic neurons during the 30-min period after heroin injection, based on data from (**b**).

**d**, Distance traveled in open field before and after morphine injection (30 mg/kg, grey dotted line) in WT (black, n = 9) and DAT-*MCU* cKO (red, n = 9) mice, shown as time course (left) and cumulative distance during 60-min post-injection period (right).

**e**, (Left) Schematic showing MCU knockout in dopaminergic neurons via bilateral AAV-TH-Cre injections into VTA of MCU floxed mice. (Right) Representative immunofluorescence images confirming knockout efficacy, showing MCU (green) and TH (red) expression in the VTA of wild-type (top) and knockout (bottom) mice.

**f**, Distance traveled in open field for $MCU^{flox/flox}$ sham (black) and $MCU^{flox/flox}$ AAV-TH-Cre (red) mice before and after heroin (left, 10 mg/kg) or cocaine (right, 20 mg/kg) injection. Grey dotted lines and black arrowheads mark injection times. Sample sizes: heroin (sham : $n = 8$; TH-Cre: $n = 7$) and cocaine (sham : $n = 8$; TH-Cre: $n = 8$).

**g**, Cumulative distance traveled within 60 minutes after heroin or cocaine injection, derived from locomotor activity data in (**d**).

**h**, CPP scores during pre-test and post-test sessions for heroin ($n = 12$ per group) and cocaine ($n = 10$ per group) in $MCU^{flox/flox}$ sham and $MCU^{flox/flox}$ AAV-TH-Cre mice .

**i**,**j**, Locomotor activity during sensitization to heroin (10 mg/kg, **i**) or cocaine (20 mg/kg, **j**) within 60 minutes post-injection. Data are shown for $MCU^{flox/flox}$ sham (n = 8 for heroin; n = 7 for cocaine) and $MCU^{flox/flox}$ AAV-TH-Cre (n = 7 per group) mice after 3 days of saline pretreatment.

All data are represented as mean ± SEM. Statistical analyses were conducted using two-sided unpaired Student's t-test (**c**, **g**) and two-way ANOVA with Bonferroni post hoc test (**d**, **f**, **h**–**j**). NS, not significant; $^{*}P < 0.05$, $^{**}P < 0.01$, $^{***}P < 0.001$.

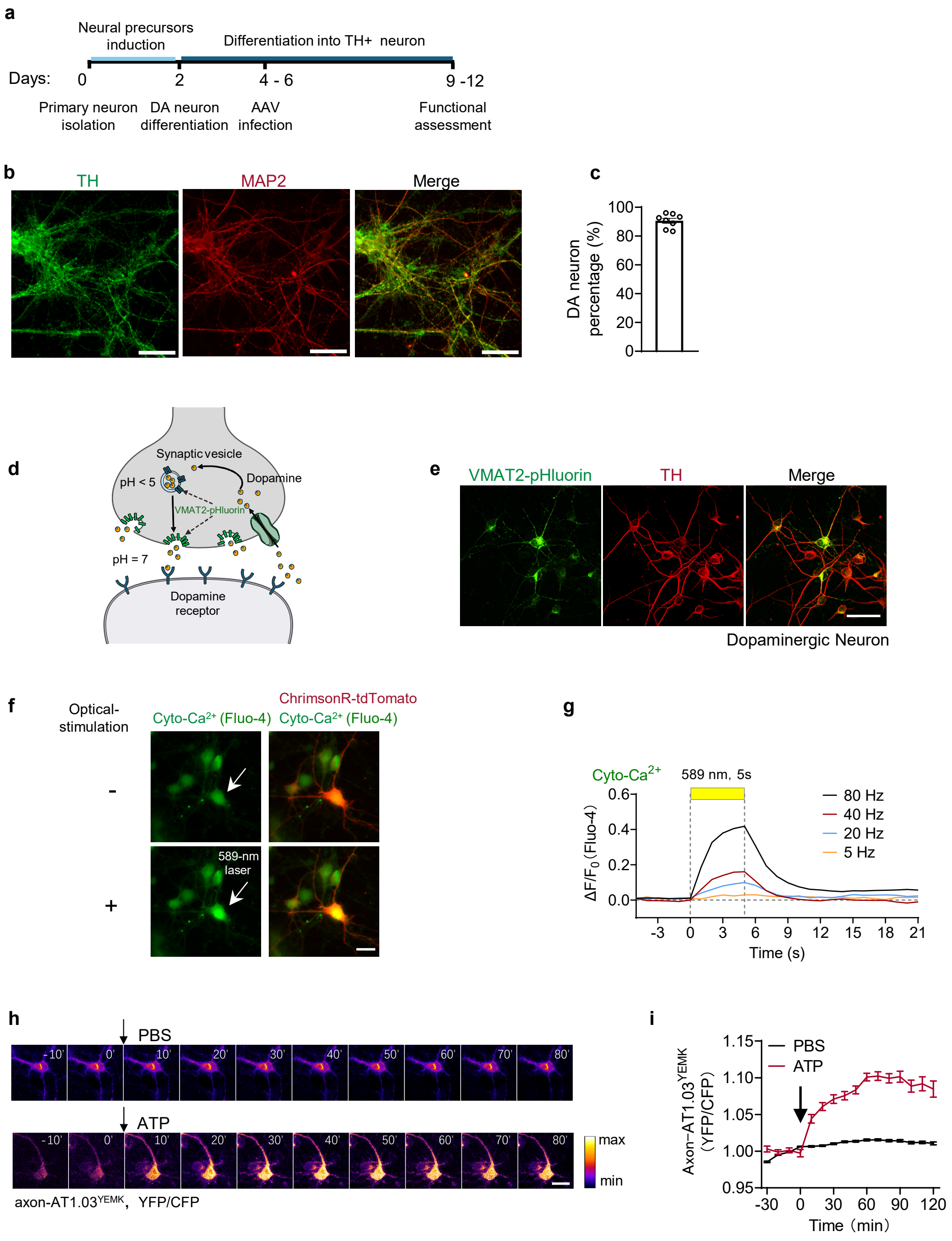


**Extended Data Figure 3. Establishment of mouse dopaminergic neuron primary cultures for *in vitro* applications.**

**a**, Schematic of the experimental timeline for differentiating primary neuron into tyrosine hydroxylase-positive ($TH^+$) dopaminergic neurons, including AAV infection.

**b,c**, Representative immunofluorescence images (**b**) and quantitative analysis (**c**) demonstrating efficient dopaminergic neuronal differentiation, with approximately 90% of MAP2-positive neurons (red) expressing tyrosine hydroxylase (TH, green). Scale bar: 50 μm.

**d**, Schematic of VMAT2-pHluorin fluorescent reporter system, designed for real-time analysis of vesicular neurotransmitter recycling dynamics.

**e**, Representative images of VMAT2-pHluorin (green) specifically expressed in dopaminergic neurons (TH, red). Scale bar: 50 μm.

**f**, Representative fluorescence images showing cytoplasmic calcium responses in ChrimsonR-tdTomato-expressing dopaminergic neurons (red) loaded with Fluo-4 AM (green), before (-) and after (+) 589-nm laser stimulation. Arrow indicates the stimulated neuron. Scale bar: 20 μm.

**g**, Representative traces of cytoplasmic calcium responses (Fluo-4 AM) in DA neurons to optogenetic stimulation at various intensities. (589 nm laser, 5 ~ 80 Hz, 5 s duration, 3 ms square wave pulses, 4 mW at the fiber tip).

**h**, Time-lapse images of axonal ATP level changes in neurons expressing axon-AT1.03$^{YEMK}$, measured by the YFP/CFP emission ratio after the adding PBS or 5 mM ATP. Scale bar: 20 μm. The arrow marks the time of PBS/ATP application.

**i**, Traces of axon ATP levels (axon-AT1.03$^{YEMK}$) following PBS ($n$ = 60 neurons) or 5 mM ATP ($n$ = 54 neurons) treatment. Arrow marks the time of PBS/ATP application.

All data are represented as mean ± SEM. Experiments were independently repeated three times.

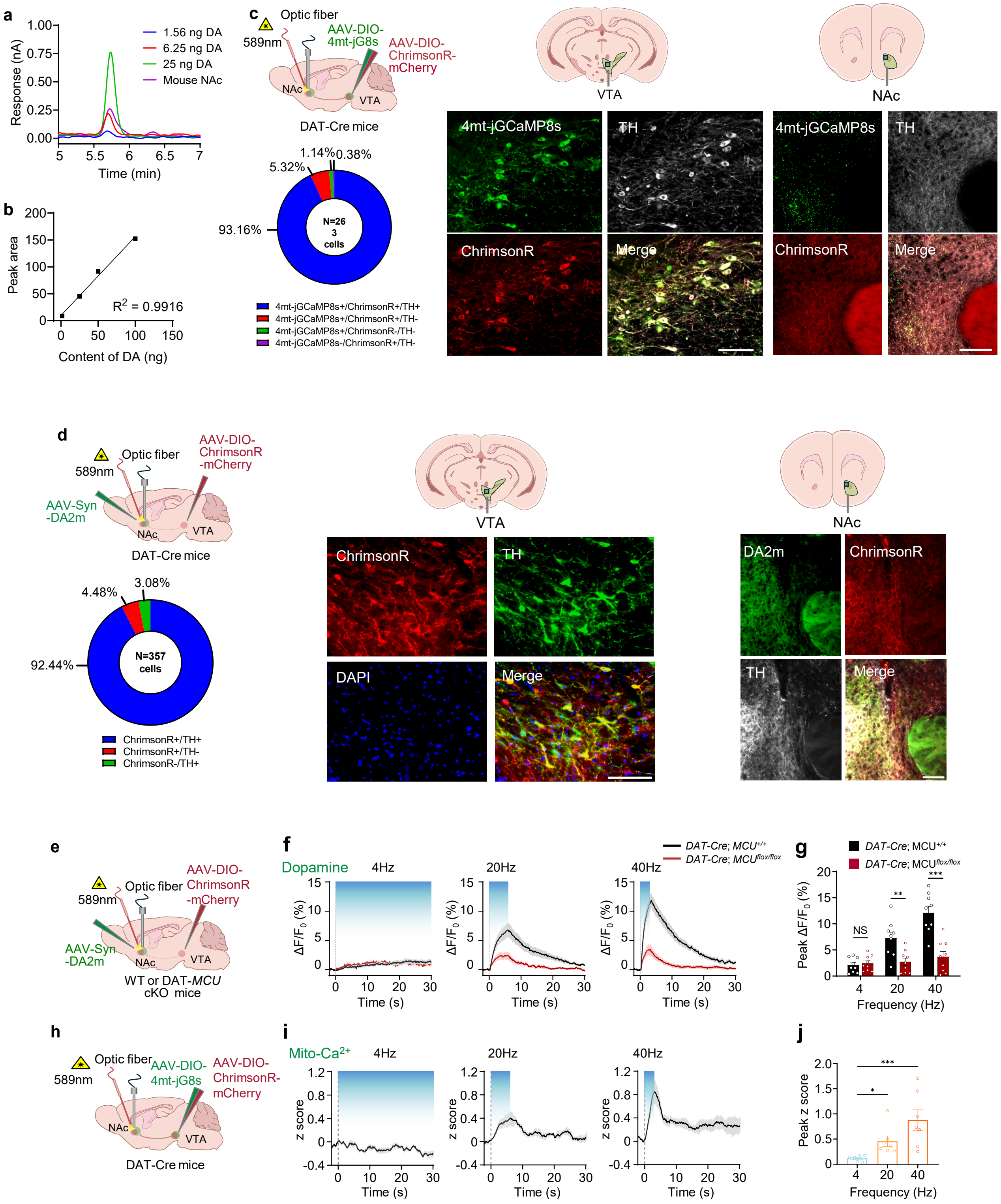


**Extended Data Figure 4. MCU knockout impairs high-intensity dopamine release without affecting basal dopamine levels.**

**a**, HPLC chromatograms of three dopamine standards and dopamine from NAc tissue sample.

**b**, Standard curve generated from the spiked dopamine concentrations.

**c**, (Left) Top: Schematic of unilateral stereotaxic injection of AAV-DIO-4mt-jGCaMP8s and AAV-DIO-ChrimsonR-mCherry into the VTA, followed by fiber implantation and optogenetic stimulation in the NAc. Bottom: High co-expression of 4mt-jGCaMP8s and ChrimsonR and their specificity to $TH^+$ neurons in the VTA. (Right) Representative images showing co-expression of ChrimsonR and 4mt-jGCaMP8s in $TH^+$ neurons within the VTA and NAc. Scale bar: 100 μm.

**d**, (Left) Top: Schematic of unilateral stereotaxic injection of AAV-DIO-4mt-jGCaMP8s into the VTA, AAV-Syn-DA2m into the NAc; and fiber implantation followed by optogenetic stimulation in the NAc. Bottom: ChrimsonR expression is highly specific to $TH^+$ neurons in the VTA. (Right) Representative images showing ChrimsonR specifically expressed in $TH^+$ neurons within the VTA and NAc. Scale bar: 100 μm.

**e**, Schematic showing fiber photometry setup for monitoring dopamine dynamics during optogenetic stimulation in WT and DAT-*MCU* cKO mice.

**f**, Traces of dopamine signals (DA2m, $\Delta F/F_0$) in the NAc before and after 120 pulses optogenetic stimulation (blue shading) with 4, 20, or 40 Hz in WT and DAT-*MCU* cKO mice ($n = 9$ mice per group).

**g**, Bar graph showing peak $\Delta F/F_0$ of the dopamine signals during optogenetic stimulation (120 pulses) with different frequencies in WT and DAT-*MCU* cKO mice, based on the data in (**f**) ($n = 9$ mice per group).

**h**, Schematic showing fiber photometry setup for monitoring mitochondrial $Ca^{2+}$ dynamics during optogenetic stimulation in DAT-Cre mice .

**i**, Traces of mitochondrial $Ca^{2+}$ dynamics (4mt-jGCaMP8, z score) in NAc dopaminergic terminals before and after 120 pulses optogenetic stimulation (blue shading) with 4, 20, or 40 Hz ($n = 7$ mice per group).

**j**, Bar graph showing peak z score of the mitochondrial $Ca^{2+}$ signals during optogenetic stimulation (120 pulses) with different frequencies, based on the data in (**i**) ($n = 7$ mice per group).

All data are presented as mean ± SEM. Statistical analyses were conducted using Pearson correlation test (**b**), two-sided unpaired Student's t-test (**g**) and Kruskal-Wallis test with a Dunn's multiple comparison test (**j**). NS, not significant; $^{*}P < 0.05$, $^{**}P < 0.01$, $^{***}P < 0.001$.

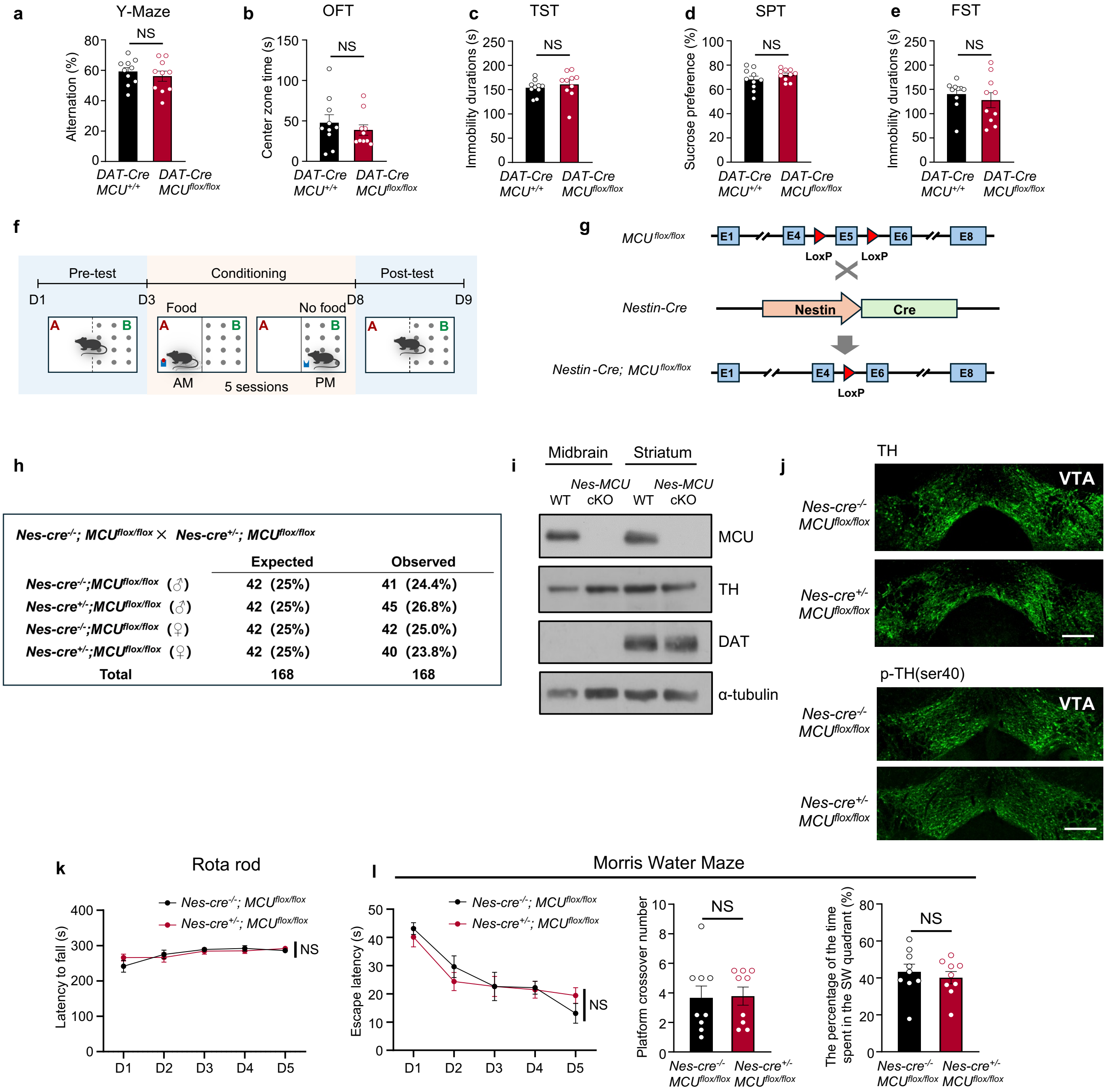


**Extended Data Figure 5. MCU knockout does not affect dopamine-related physiological functions in mice.**

**a–e**, Behavioral tests comparing *DAT-Cre; MCU$^{flox/flox}$* mice ($n$ = 10) and *DAT-Cre; MCU$^{+/+}$* mice ($n$ = 10) reveal no significant differences in: **a**. Alternation behavior in the Y maze test. **b.** Time spent in the center zone in the open-field test (OFT). **c.** Duration of immobility in the tail suspension test (TST). **d.** Sucrose preference in the sucrose preference test (SPT). **e.** Duration of immobility in the forced swimming test (FST).

**f**, Experimental timeline of food-conditioned place preference (CPP) corresponding to Fig. 5q.

**g**, Genetic strategy for whole-brain MCU knockout using Nes-Cre mice.

**h**, Offspring frequencies from *Nes-Cre$^{-/-}$; MCU$^{flox/flox}$* crosses with *Nes-Cre$^{+/-}$; MCU$^{flox/flox}$* mice were consistent with Mendelian ratios, with no deviation in genotype distribution or sex ratio, suggesting no fetal mortality due to whole-brain MCU deletion.

**i**, Western blot analysis of MCU, TH, and DAT protein expression in midbrain and striatum of *Nes-Cre$^{+/-}$; MCU$^{flox/flox}$* mice and *Nes-Cre$^{-/-}$; MCU$^{flox/flox}$*.

**j**, Expression levels of tyrosine hydroxylase (TH) and phosphorylated TH (p-TH) in the VTA were comparable between *Nes-Cre*$^{+/-}$*; MCU*$^{flox/flox}$ mice and *Nes-Cre*$^{-/-}$*; MCU*$^{flox/flox}$ mice, suggesting that MCU deletion does not affect dopaminergic neurons. Scale bar: 200 µm.

**k**, Latency to fall on the rota-rod test, measuring motor coordination, was equivalent between *Nes-Cre*$^{-/-}$*; MCU*$^{flox/flox}$ and *Nes-Cre*$^{+/-}$*; MCU*$^{flox/flox}$ mice($n$ = 9 per group), suggesting no impairment in motor function.

**l**, Morris water maze test results show no differences between *Nes-Cre*$^{-/-}$*; MCU*$^{flox/flox}$ and *Nes-Cre*$^{+/-}$*; MCU*$^{flox/flox}$ mice ($n$ = 9 per group) in: (Left) Learning curve show latency to find the hidden platform over a 5-day training period. (Middle) Number of platform site crossings during the probe test. (Right) Percentage of time spent in each quadrant during the probe test with the platform removed, indicating that MCU deletion does not affect learning or memory.

All data are presented as mean ± SEM. Statistical analyses were conducted using two-sided unpaired Student's t-test (**a**–**d**, **l**-right), Mann Whitney test (**e**, **l**-center), and two-way ANOVA with post-hoc Bonferroni (**k**, **l**-left). NS, not significant; $^{*}P < 0.05$, $^{**}P < 0.01$, $^{***}P < 0.001$.

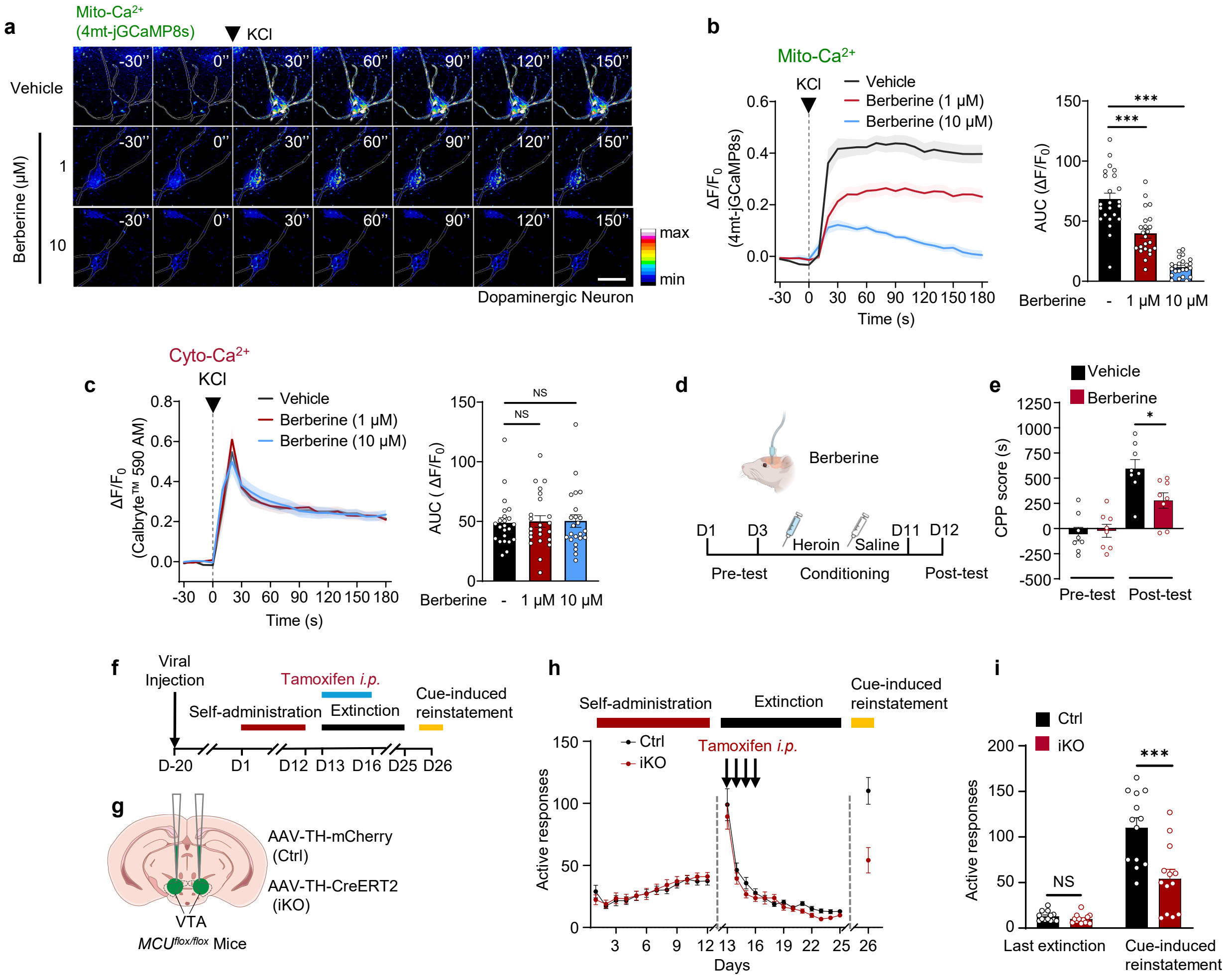


**Extended Data Figure 6. Pharmacological and genetic inhibition of MCU suppresses drug reward and drug-seeking behaviors.**

**a**, Pseudocolored time-lapse fluorescence images of dopaminergic neurons expressing 4mt-jGCaMP8s after KCl stimulation (75 mM). Pre-treatments included vehicle (top), 1 μM berberine (middle), and 10 μM berberine (bottom). The arrow marks the time of KCl stimulation. Scale bar: 50 μm.

**b**,**c**, Traces (left) and area under the curve (AUC) values (right) of mitochondrial $Ca^{2+}$ (**b**) and cytoplasmic $Ca^{2+}$ (**c**) responses to KCl stimulation. Data shown for vehicle-treated (black), 1 μM berberine-treated (red), and 10 μM berberine-treated (blue) groups ($n$ = 23 neurons per group). The dashed arrow indicates the KCl stimulation time.

**d**, Schematic of the experimental design for heroin-induced conditioned place preference (CPP), including timing for intra-NAc microinjections (0.3 μg per side berberine administered 30 minutes before each heroin injection during training).

**e**, Berberine inhibits the acquisition of heroin-induced CPP, as shown by reduced CPP scores in post-test compared to the vehicle group ($n$ = 8 per group).

**f**, Experimental timeline of Cre-ERT2 mediated inducible MUC conditional knockout in cocaine self-administration paradigm, including AAV-TH-CreERT2 injection, cocaine self-administration training, extinction, tamoxifen-induced recombination and cue-induced seeking test.

**g**, Schematic illustration of inducible conditional MCU knockout strategy in VTA dopaminergic neurons via bilateral stereotaxic delivery of AAV-DIO-TH-mCherry or AAV-DIO-TH-CreERT2 into the VTA of the $MCU^{flox/flox}$ mice.

**h**, Active nose-poke responses during cocaine self-administration training, extinction, and cue-induced reinstatement test in Ctrl and iKO mice (n = 13 per group). Arrows indicate daily intraperitoneal administration of tamoxifen for 4 consecutive days starting from the first day of extinction.

**i**, Active nose-poke responses between Ctrl and iKO during the final extinction session and cue-induced reinstatement test of cocaine self-administration.

All data are presented as mean ± SEM. Statistical analyses were conducted using one-way ANOVA with Bonferroni multiple comparison test (**b**-right), Kruskal-Wallis test with a Dunn's multiple comparison test (**c**-right) and two-way ANOVA with Bonferroni post hoc test (**e**, **h**, **i**). NS, not significant; $^{*}P < 0.05$, $^{**}P < 0.01$, $^{***}P < 0.001$.

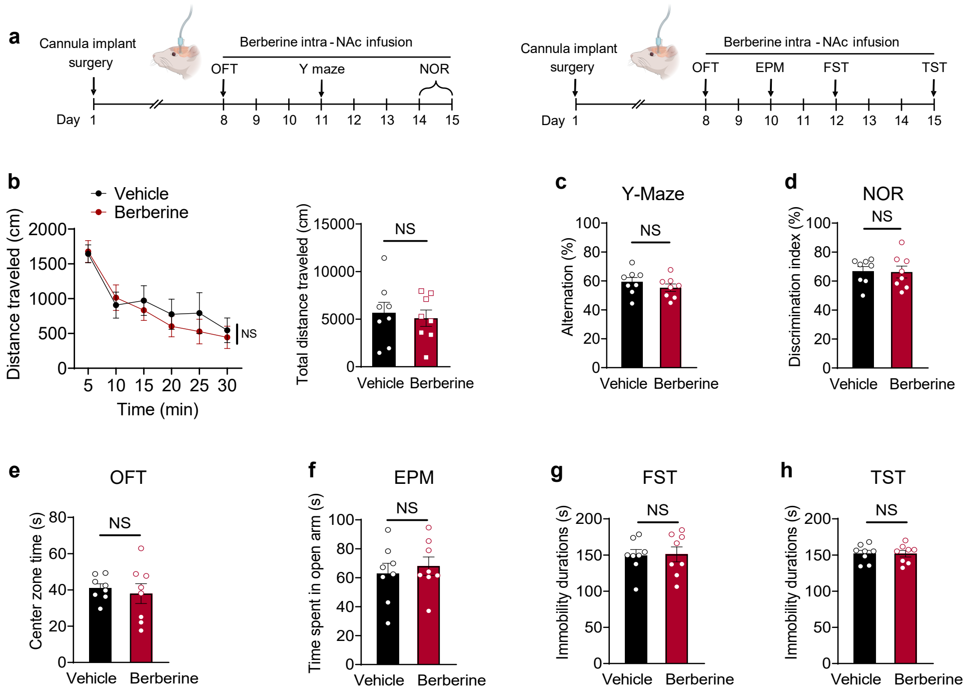


**Extended Data Figure 7. Berberine does not affect basal dopamine-related physiological functions in mice.**

**a**, Schematic of the experiment timeline, including intra-NAc microinjection of berberine and subsequent behavioral analyses.

**b**, Locomotor activity assessed via open-field test after intra-NAc infusion of berberine or vehicle. Left: distance traveled every 5 minutes during a 30-minute test. Right: total distance traveled by each group (n = 8 per group).

**c**,**d**, No significant differences in alternation behavior in the Y maze test (**c**) or the discrimination index in the novel object recognition test (NOR) (**d**) between vehicle- and berberine-treated mice, indicating that berberine does not impair spatial working memory or object recognition memory ($n$ = 8 per group).

**e**,**f**, Time spent in the center zone of the open-field test (OFT) (**e**) and the open arms of the elevated plus maze (EPM) (**f**) showed not significant differences between groups (n = 8 per group), indicating no effect on anxiety-like behavior.

**g**,**h**, Immobility durations in forced swimming test (FST) (**g**) and tail suspension test (TST) (**h**) were comparable between berberine- and vehicle- treated groups($n$ = 8 per group), suggesting no effect on depressive-like behavior.

All data are presented as mean ± SEM. Statistical analyses were conducted using two-sided unpaired Student's t-test (**b**-right, **c**–**h**), and two-way ANOVA with Bonferroni post hoc test (**b**-left). NS, not significant.
